# Directed evolution of multimeric proteins is enabled by dual-compensatory gene duplication

**DOI:** 10.64898/2026.01.12.698938

**Authors:** Rezwan Siddiquee, Felicia Lie, Taylor N. Szyszka, Alex Loustau, Michael P. Andreas, Tobias W. Giessen, Yu Heng Lau

## Abstract

Gene duplication has played a critical role in the evolutionary history of proteins, enabling complex multimers to emerge from simpler precursors. Yet in protein engineering, current methods for directed evolution do not exploit gene duplication, hampering access to the vast array of diverse variants that are only enriched in the presence of a wild-type copy. We establish a directed evolution strategy for multimeric proteins that harnesses gene duplication to compensate for metabolic burden and self-assembly fitness, allowing previously inaccessible variants to be enriched. Starting from a homomeric 240-mer capsid, gene duplication enables selection of both extreme homomeric variants and obligate heteromers. This strategy significantly expands engineering access to diverse high-performing variants, while also supporting a plausible model for evolutionary diversification of higher-order multimers in nature.

## Main Text

Across the myriad of diverse proteins that have evolved in nature, multimeric structures are highly prevalent (*1–3*). Many multimers are thought to have arisen through the process of gene duplication (*4–6*), where a second copy of the encoding gene provides capacity for diversification (*7*). Although many models have been proposed over the past 50+ years to explain the role of gene duplication in ancestral evolution (*4, 6, 8–11*), recent studies are only now reporting experiments which demonstrate the range of outcomes that can emerge at the protein level (*12, 13*). Gene duplication followed by simple mutations can lead to the formation of higher-order multimers and the transition from homomers to heteromers (*12, 14–17*), even in the absence of a strong selection pressure or adaptive gain (*18, 19*). In other instances, duplication may be accompanied by paralog interference that can encourage additional complexity or preservation of genetic redundancy (*20, 21*). While not always leading to enhanced functionality, duplication facilitates opportunities for broader exploration of the protein fitness landscape.

In the field of protein engineering, basic evolutionary principles have been co-opted for the selection of variants with desirable traits, yet typical directed evolution campaigns do not exploit gene duplication (*22, 23*). Unlike ancestral protein evolution events, laboratory directed evolution requires a robust genetic design such that when a selection pressure is applied, high-fitness variants that exhibit the desired performance characteristics are enriched. Undesirable selection outcomes can occur when the genetic design does not adequately account for the multitude of confounding factors that contribute to overall fitness, such as metabolic burden, expression level, solubility, and stability (*24–26*). As variants are typically generated from a single genetic copy, these confounders can severely limit the diversity of the accessible protein sequence landscape. This effect is exacerbated for multimeric proteins when the impact of a deleterious mutation is amplified by the repeated extensive interactions between adjacent subunits.

In this study, we implement gene duplication in the directed evolution of protein homomers (Fig. 1A), compensating for confounding factors to enable selection of more diverse and otherwise inaccessible variants, including non-native obligate heteromers with adaptive function. Our model system is the encapsulin protein from *Quasibacillus thermotolerans* (QtEnc), which self-assembles to form a homomeric 240-mer cage structure that shares homology with the widespread HK97 family of viral capsids (*27*). Encapsulin systems have attracted significant interest in biotechnology as scaffolds for antigen display (*28*), drug delivery (*29*), microscopy (*30, 31*), and enzyme encapsulation (*32, 33*), where the ability to evolve novel variants would be a powerful method for engineering and optimizing designs (*34*). Here, we develop a directed evolution system which selects for encapsulin assemblies with increased porosity (*35, 36*), where gene duplication is an essential feature designed to compensate for confounding changes in both assembly fitness and metabolic burden.

**Fig. 1.**
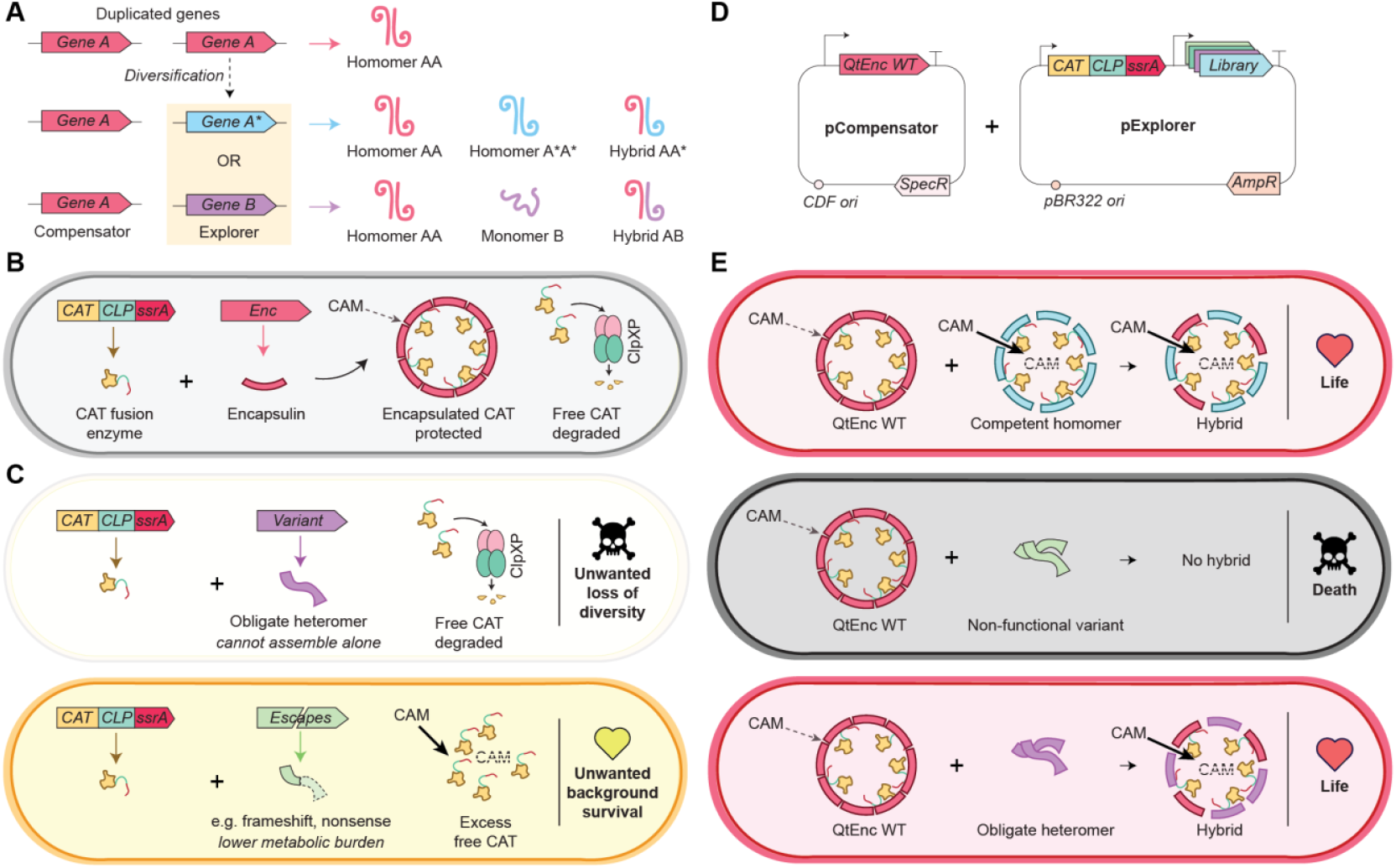
A gene duplication circuit enables divergent directed evolution of higher-order multimeric proteins. (**A**) Schematic depicting how gene duplication can enable divergent evolution from homomeric to hybrid heteromeric complexes. The Compensator copy (Gene A) encodes for the wild-type protein, while the Explorer copy encodes for different variants (Gene A* or Gene B). Some variants can generate functional homomers (A*A*) and heteromers (AA*), while others can only form heteromers (AB) that are only selectable in the presence of the Compensator. ( **B**) Genetic circuit for life-death selection of encapsulin variants in *E. coli*. The enzyme chloramphenicol acetyltransferase (CAT) bearing an ssrA tag is degraded by ClpXP unless sequestered within an encapsulin via its cargo loading peptide (CLP). Encapsulation protects the enzyme but confers limited chloramphenicol (CAM) resistance due to the encapsulin serving as a semi-permeable barrier to substrate access. (**C**) Schematic showing two unwanted scenarios from selection. Obligate heteromeric variants that function only as hybrids with WT do not survive, leading to loss of diversity. Non-functional (escape) variants with lower metabolic burden can result in high background survival. ( **D**) Gene duplication strategy implemented on a two-plasmid circuit. pExplorer expresses the encapsulin variant library, while pCompensator expresses an invariant copy (QtEnc WT) to compensate for any changes in metabolic burden and assembly fitness. (**E**) Three predominant outcomes of selection are anticipated: survival of competent homomeric variants that enhance overall CAT activity, death arising from completely non-functional variants, and obligate heteromers that can enhance CAT activity upon hybridization with QtEnc WT.

## Results

### Design of a gene duplication circuit to compensate for metabolic burden and assembly fitness

We designed a life-death selection strategy for the directed evolution of encapsulins (Fig. 1B-E), where survival is based on encapsulation of the antibiotic resistance enzyme chloramphenicol acetyltransferase (CAT). At the C-terminus of CAT, we appended the native cargo-loading peptide (CLP) of QtEnc to load CAT into encapsulins, as well as an ssrA degron tag to facilitate degradation of any excess unencapsulated CAT by the endogenous ClpXP protease (Fig. 1B). QtEnc variants that enhance overall CAT activity in the expression host were expected to promote survival. This includes variants that provide encapsulated CAT with increased substrate access, as wild-type QtEnc (WT) has relatively narrow pores (2-7 Å) (*27*), along with variants that enhance the stability of encapsulated CAT. Meanwhile, variants that lose the ability to assemble were not intended to survive, as they should not protect CAT from proteolytic degradation.

We envisaged two confounding issues that could arise when conducting directed evolution on a single copy of the encapsulin gene (Fig. 1C). There may be variants of interest that are only functional as heteromeric hybrid assemblies and do not confer survival as a single copy, leading to a loss of accessible sequence diversity. In addition, non-functional encapsulin variants that incur a lower metabolic burden than the WT could lead to increased CAT expression that saturates the degradation machinery, ultimately compromising the selection due to false-positive background survival.

To avoid these two unwanted scenarios, we designed a two-plasmid genetic circuit for gene duplication, where ‘pCompensator’ expresses an invariant copy of QtEnc WT and ‘pExplorer’ contains the variant library (Fig. 1D). The inclusion of QtEnc WT allows obligate heteromeric variants to form hybrid assemblies, thus expanding the accessible landscape of sequence variants. At the same time, QtEnc WT provides a baseline level of metabolic burden and CAT encapsulation, even in the presence of non-functional variants with lower burden, thus countering false-positive survival. Three predominant selection outcomes were anticipated (Fig. 1E). First, if the variant forms a competent assembly that enhances overall CAT activity, the host survives. Second, if the variant cannot form any competent assemblies, CAT is only encapsulated in QtEnc WT and has insufficient activity to confer survival to the host. Third, if the variant is an obligate heteromer that can only assemble with QtEnc WT, the host survives in cases where the resulting hybrid assembly increases overall CAT activity.

### Directed evolution with gene duplication selects for extreme variants that enhance survival

Our library of encapsulin variants featured deletions at the vertex-lining loop of the A-domain of QtEnc (residues 190-211, Fig. 2A) to generate larger pores, a desirable trait for enhancing enzyme substrate access when encapsulins are used in nanoreactor applications (*37–39*). This region lines the pentameric and hexameric vertices of the icosahedral assembly (*27*), where the WT has narrow ∼2 Å diameter pores at the pentameric vertices and closed hexameric vertices. While prior studies have explored a handful of small rationally-designed loop deletions in encapsulins from other species (*35, 37, 38, 40*), extreme deletions (>7 residues) have not been reported due to the perceived risk of completely destabilizing the assembly.

**Fig. 2.**
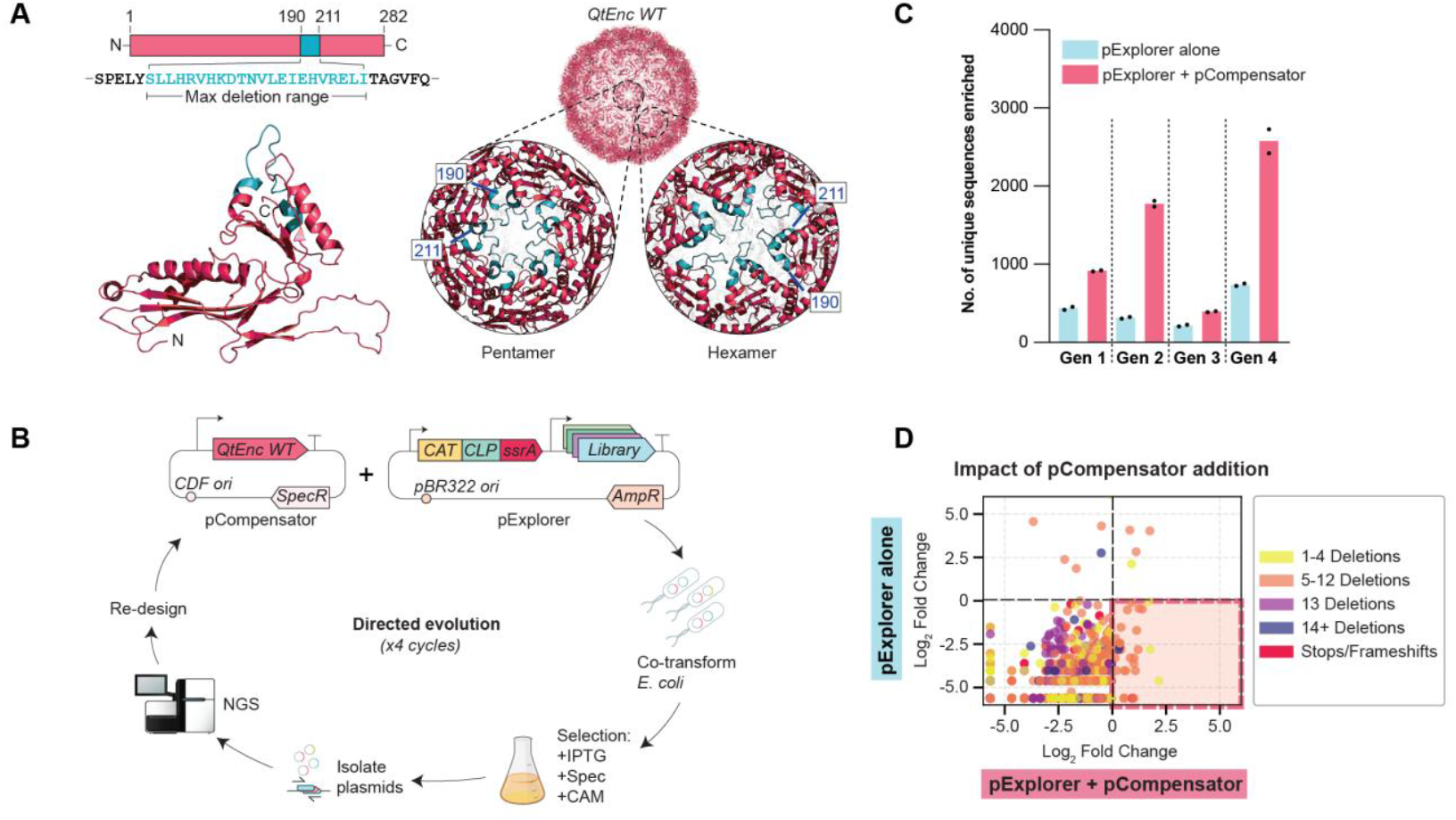
Directed evolution with compensatory gene duplication expands the accessible landscape of sequence variants. (**A**) Structures of QtEnc monomer and assembly, where the blue region indicates the maximum deletion range in the variant libraries. Deletions flank the pentameric and hexameric vertices of the icosahedral assembly. (**B**) Directed evolution workflow with four iterative rounds of selection and subsequent library re-design. (**C**) With pCompensator included, more sequence variants are enriched by directed evolution. (**D**) Comparison of fold change for each unique sequence variant with and without the inclusion of pCompensator. Plot shows data for Gen 4 selection with 15 µg/mL CAM. Sequences in the lower right quadrant (highlighted in pink) were only enriched with pCompensator, representing a unique set of variants that is made accessible by compensatory gene duplication. Each encapsulin variant is represented as a single dot, whose x-coordinate reflects fold change over input library after selection when pCompensator is included, and y-coordinate reflects fold change when pCompensator is absent. Dot points are colored according to number of deletions in particular variant.

We performed directed evolution with four generations of encapsulin libraries (Fig. 2B and Fig. S1-4), finding that stringent chloramphenicol selection with the inclusion of gene duplication provides reliable access to extreme deletion variants that would otherwise have been purged (Fig. 2C-D). After initial optimization of CAT promoter strength, we identified selection conditions where strains expressing QtEnc WT displayed minimal growth as a baseline for stringent selection (Fig. S5). Upon conducting directed evolution, sequencing analysis of surviving variants across all four generations showed that the inclusion of pCompensator significantly expanded the number and diversity of enriched sequences compared to traditional single-copy selection (Fig. 2C-D), while also reducing the frequency of unwanted false-positive frameshifts and nonsense mutations that would otherwise have been enriched in some rounds of selection (Fig. S6).

The ‘Left/Right’ library (Gen 1) featured incremental unidirectional deletions extending from the pore region toward either the N-terminus (‘Left’: residues 203-190) or C-terminus (‘Right’: residues 197-211). Each deletion variant included a random residue substitution encoded by an NNK codon adjacent to the deletion site to introduce local diversity (Fig. 3A and Fig. S1). Selection was performed with chloramphenicol as selection pressure and spectinomycin to retain the compensatory copy of QtEnc WT. Surviving sequences were compared against the input library with no antibiotic selection pressure. Results showed that while the input library contained diverse variants with up to 15 residues deleted, the selected population was dominated by conservative variants with 1-4 residues deleted, while larger deletions were purged (Fig. 3A). This result suggested that simple loop excision did not provide sufficient flexibility to maintain structural integrity at the pore-lining region.

**Fig. 3.**
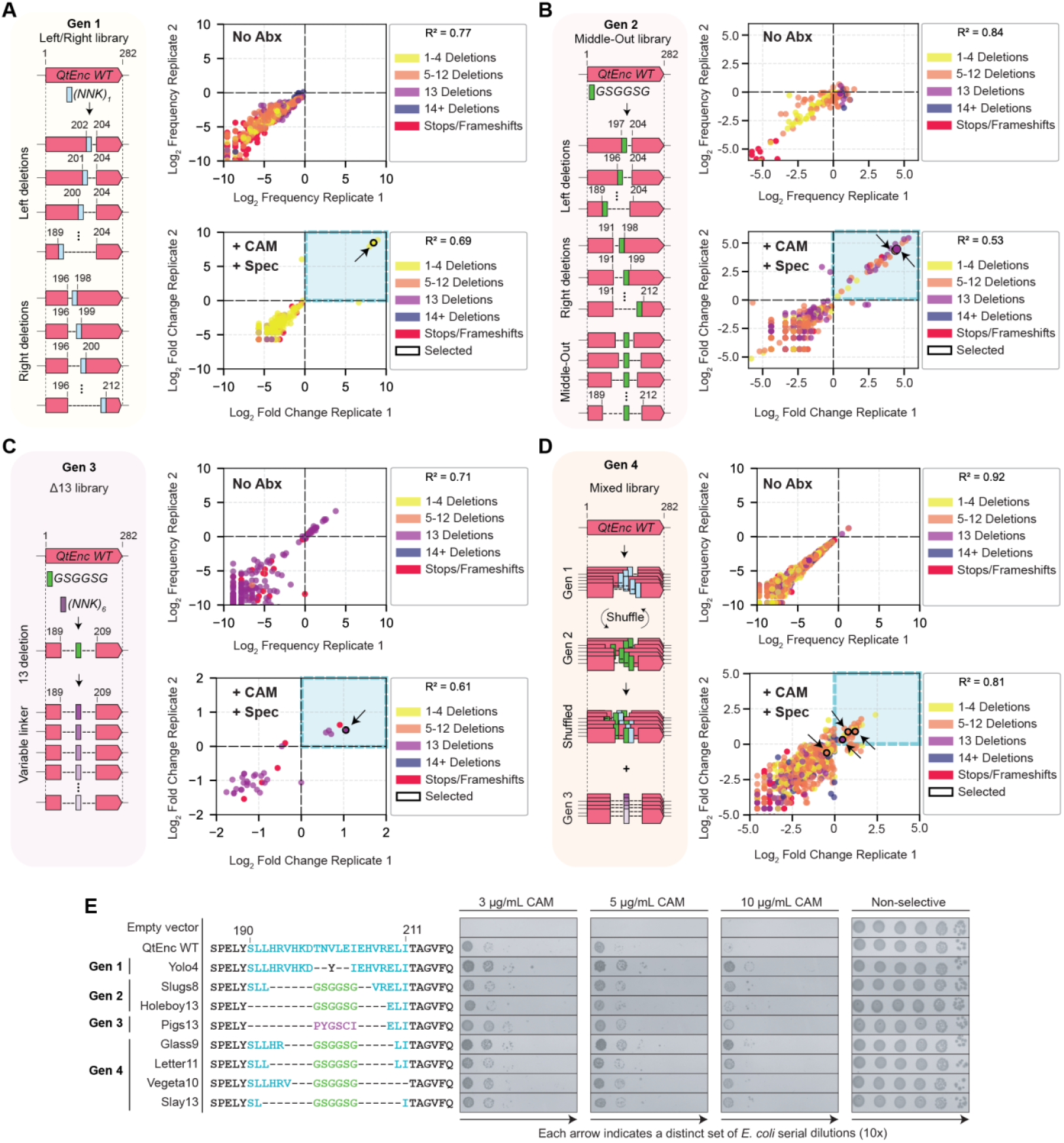
Directed evolution selects for diverse deletion variants that enhance survival. (**A-D**) Four generations of variant libraries in pExplorer - Gen 1: Left/Right, Gen 2: Middle-Out, Gen 3: Δ13, and Gen 4: Mixed. Log_2_ frequency plots (as a proportion of total reads) are shown for experimental replicates of input library without selection antibiotics (No Abx), i.e. no chloramphenicol (CAM) and no spectinomycin (Spec), while Log _2_ plots of fold change over input library are shown for experimental replicates with selection (CAM and Spec). R^2^ values indicate the degree of linear correlation between replicates. Each encapsulin variant is represented as a single dot, colored according to the number of deletions in each variant sequence. Selected variants for further study have a black outline and are indicated with an arrow. (**E**) Selected QtEnc variants in pExplorer spotted on selective plates (3-10 µg/mL CAM, 0.1 mM IPTG) show approximately an order of magnitude greater survival than WT. Residues within the maximum deletion range are shown in blue, while GSGGSG linkers are in green, and the random linker is in purple.

To restore structural continuity, we designed a ‘Middle-Out’ library (Gen 2) featuring a flexible glycine-serine linker (GSGGSG) to bridge the flanking helices, with a theoretical maximum deletion size of 17 residues (Fig. 3B and Fig. S2). Sequencing of the library after selection revealed deletions of 4-13 residues, with the 13 residue deletions most strongly selected for, while no deletions of >14 residues were observed (Fig. 3B). This result suggests that deletion of 13 residues is the physical limit achievable with this linker strategy, validating that structural continuity is the primary constraint.

Exploring the sequence space of the most extreme deletion variants further, we constructed the ‘Δ13’ library (Gen 3) by replacing the defined glycine-serine linker from the Gen 2 selection results with six random residues encoded by NNK codons (Fig. 3C and Fig. S3). We expected that these large deletions would cause most variants to be assembly-incompetent on their own, while gene duplication should rescue variants that can still form competent hybrid assemblies with QtEnc WT, thus selecting for obligate heteromers. Sequencing data showed that the selection enriched for additional distinct sequences (Fig. 3C), while critically, omission of pCompensator in a negative control selection resulted in populations with overrepresentation of false-positive stop codons and frameshifts (Fig. S6).

To maximize library diversity in the final design iteration, we shuffled the Left/Right and Middle-Out libraries together using the StEP method (*41*), then added the Δ13 library (Fig. 3D, Fig. S4 and Fig. S7). This ‘Mixed’ library (Gen 4) was subjected to stringent selection to isolate the highest-performing variants at increasing chloramphenicol concentrations.

To validate the sequencing results from our directed evolution campaign, we selected representative enriched variants across all four generations for further study (Fig. 3E). These shortlisted variants were cloned individually into the pExplorer format (i.e. co-expressed with CAT-CLP-ssrA), with growth assayed by spotting serial dilutions onto plates with different concentrations of chloramphenicol. All variants conferred greater chloramphenicol resistance than QtEnc WT, confirming that sequence enrichment was driven by enhanced survivability as intended.

### Extreme deletion variants can assemble into stable homomeric cages

From the shortlist of encapsulin variants, we chose two top-performing variants enriched from the Gen 4 library, ‘Glass9’ (9-residue deletion) and ‘Letter11’ (11-residue deletion), to characterize their ability to assemble independently of QtEnc WT. Both variants were able to form stable, monodisperse homomeric assemblies when purified from *E. coli* (Fig. 4A). Cryo-EM reconstruction of Glass9 confirmed formation of the canonical T=4 architecture, albeit with significantly enlarged pores of 21 Å diameter at the pentameric vertices and elongated 12 × 35 Å pores at the hexameric vertices (Fig. 4B-C and Fig. S8). Unexpectedly, Cryo-EM analysis of Letter11 revealed structural plasticity. The population was dimorphic (Fig. 4B), containing the expected T=4 assemblies (∼70%) as well as smaller 180-subunit T=3 assemblies (∼30%) (Fig. 4C and Fig. S9). This inadvertent access to a latent geometry is consistent with the structural polymorphism observed in viral capsids, virus-like particles, and some other encapsulins (*42*), but had not been reported for QtEnc, demonstrating that compensatory evolution can relax geometrical constraints to access novel architectures.

**Fig. 4.**
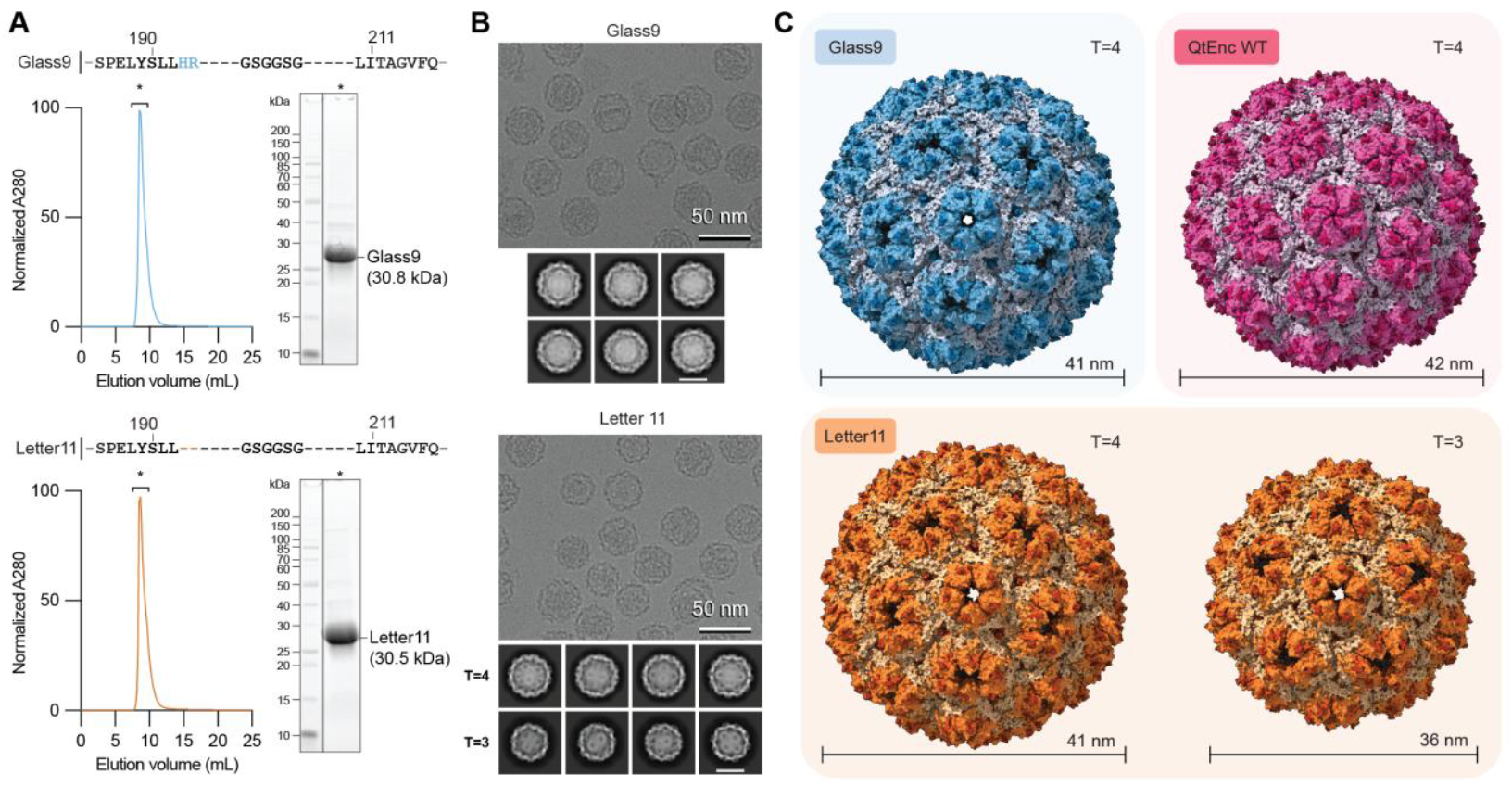
Cryo-EM structures of selected homomeric variants. (**A**) Size-exclusion chromatography on a Superose 6 Increase 10/300 GL column shows that variants Glass9 and Letter11 elute in the expected range for high molecular weight complexes, along with SDS-PAGE analysis to demonstrate purity of the elution peaks (marked with asterisks). (**B**) Representative cryo-EM micrographs indicate sample homogeneity for Glass9 and dimorphism for Letter11. Representative 2D class averages are shown below each micrograph (2D class average scale bars = 24 nm). (**C**) Structural models of Glass9 and Letter11 homomeric capsids. Glass9 (PDB 9ZVV) shares the same T=4 architecture as QtEnc WT (PDB 6NJ8), but is more porous. Letter11 forms both T=4 (PDB 9ZW3) and T=3 (PDB 9ZW4) assembly states.

### Gene duplication enables positive selection of obligate heteromers

We investigated whether our gene duplication strategy could generate obligate heteromeric variants that require hybridization with QtEnc WT to assemble. To achieve precise control of bulk stoichiometry in hybridization experiments, we applied our *in vitro* method for encapsulin assembly (*43*), triggered by cleavage of a N-terminally fused steric blockade (here using maltose-binding protein, MBP) with TEV protease (Fig. 5A). We first validated that *in vitro* assembly recapitulated the behavior observed in standard *E. coli* expression by analytical size-exclusion chromatography (SEC), with the expected high molecular weight peak observed for QtEnc WT, Glass9, and Letter11 (Fig. 5B).

**Fig. 5.**
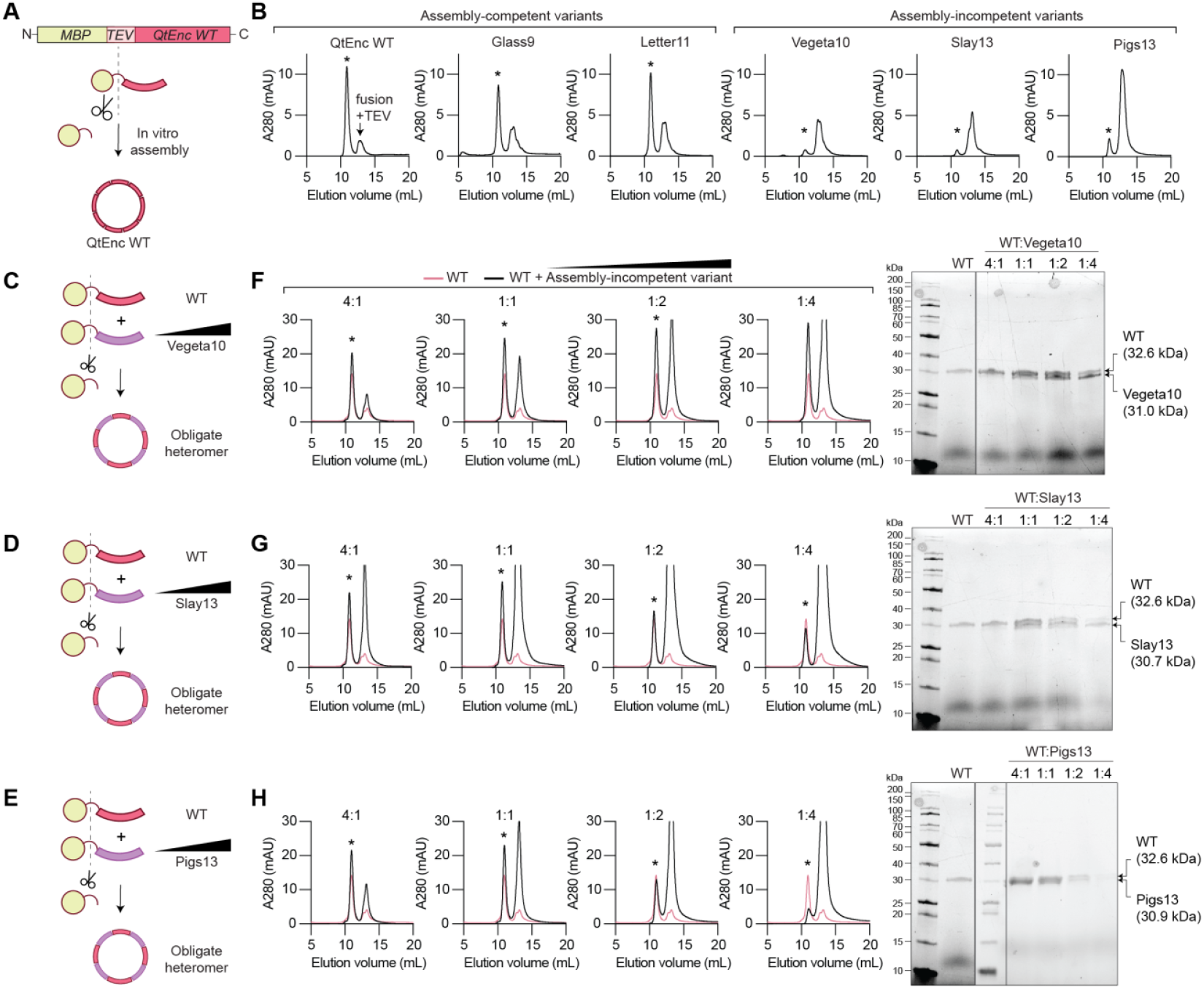
Assembly-incompetent variants can form obligate heteromeric hybrid cages. (**A**) Triggered *in vitro* assembly enables precise control of bulk stoichiometry. Cleavage of an N-terminal MBP tag with TEV protease induces spontaneous assembly. (**B**) Size-exclusion chromatography on a Bio SEC-5 2000 Å HPLC column confirms the expected elution peak (marked with asterisk) for QtEnc WT and assembly-competent variants (Glass9, Letter11), while assembly-incompetent variants (Vegeta10, Slay13, Pigs13) do not show a significant peak at the same elution volume. (**C**-**E**) Titration of pre-assembly monomers of QtEnc WT (20 µM) with assembly-incompetent variants (5, 20, 40, 80 µM), followed by *in vitro* assembly. (**F**-**H**) Vegeta10 shows increased intensity for the assembly peak relative to the WT-only chromatograph (overlaid in pink), indicating formation of hybrid assemblies. Meanwhile, dominant-negative variants Slay13 and Pigs13 form hybrid assemblies at lower ratios, but disrupt assembly at higher ratios. SDS-PAGE analysis of the elution peak for encapsulin assembly confirms the presence of both WT and variant monomers in all cases, and provides further evidence of the dominant-negative effect. The same data for the WT sample is shown in all chromatograms and gels to aid visual comparison.

The three remaining variants shortlisted from Gen 3 and Gen 4 (Vegeta10, Slay13, Pigs13) were found to be assembly-incompetent by *in vitro* assembly and analytical SEC (Fig. 5B). We then titrated these assembly-incompetent variants with QtEnc WT in their pre-assembly monomeric fusion states (Fig. 5C-E), finding that upon triggering assembly, all three variants could successfully co-assemble into hybrid cages (Fig. 5F-H). Notably, while all three variants efficiently formed hybrid assemblies at a 1:1 ratio, divergent behavior was observed at other ratios. Hybrid assembly with Vegeta10 was permissive at all ratios tested (Fig. 5F). Meanwhile, Slay13 and Pigs13 exhibited a dominant-negative effect, effectively poisoning assembly when added in higher proportions (Fig. 5G-H). This result confirms that our gene duplication strategy enabled enrichment of divergent variants that function solely within a heteromeric context, where the WT is responsible for enhancing assembly fitness while the variant is responsible for increasing the porosity, bearing similarity to sub-functionalization events that have occurred in ancestral protein evolution (*18*).

### Assembly-competent variants can form hybrid assemblies with obligate heteromeric variants

We further investigated whether assembly-incompetent variants could also form hybrids with the competent variants. Titrating the assembly-incompetent Vegeta10 and Pigs13 monomers with the assembly-competent Glass9 and Letter11 variants gave similar results to the WT *in vitro* hybridization experiments. SEC analysis showed increased A280 absorbance in the expected elution peak (Fig. S10), indicating successful heteromerization between evolved encapsulin variants, forming assemblies of similar size to QtEnc WT as determined by dynamic light scattering (Fig. S11). SDS-PAGE analysis displayed a single band with increased intensity, consistent with the near-identical molecular weights of the hybridized deletion variants (Fig. S10).

Although structural models of the hybrid assemblies could not be obtained by cryo-EM due to insufficient resolvable differences between the hybridized subunits, the data was consistent with the presence of both variant subunits in stochastically assembled particles (Fig. S12-17). Specifically, in symmetry-averaged hybrid shells, an increase in local resolution and decrease in map quality around the pentameric and hexameric pores could be observed (Fig. S15), consistent with increased variability due to stochastic occupancy. Asymmetric reconstructions of mixed shells exhibited uncharacteristically irregular pore regions compared to one-component shells (Fig. S16), further supported by high variability observed in 3D classifications of the pore regions (Fig. S17). Overall, these variant-variant hybrids showcase how directed evolution with compensatory gene duplication unlocks a ‘cryptic’ fitness landscape with variants that are only functional as heteromeric hybrids, allowing the construction of extreme variant-variant hybrid architectures that are not accessible by traditional directed evolution and homomeric assembly.

## Discussion

Our experiments show that incorporating gene duplication into a genetic circuit design enables the divergent directed evolution of multimeric structures and their function. The strategy of compensating for confounding changes in fitness associated with burden and assembly represents a generalizable framework for engineering complex molecular assemblies with greater diversity. Using QtEnc as a model homomer, we have gained unprecedented access to extreme encapsulin variants (13-residue deletions) that enhance survival based on encapsulated enzyme function, unveiled latent structural polymorphisms (T=3), and generated stable obligate heteromers. The latter observation is particularly striking, as the transition from homomer to heteromer is not seen in traditional laboratory directed evolution campaigns for biotechnological applications. While increased complexity does not always guarantee adaptive functionality, access to a broader sequence landscape increases the likelihood of unlocking new structures such as variant-variant hybrids with improved properties that can be functionally selected for.

The strong dependence of successful selection on gene duplication is likely due in part to the strong selection pressure applied in our experiments, as is the case for most life-death directed evolution campaigns. False positive variants can be strongly enriched due to uncontrolled confounding variables that enable survival. Compensating for the most important confounders allows the full potential of directed evolution to be harnessed, as well as providing greater clarity on the boundaries for obtaining functional sequences. Indeed, the most extreme variants that we were able to enrich (Pigs13 and Slay13) required the most compensation from WT to be selected for as heteromers, while slightly more conservative variants (Glass9 and Letter11) could function independently.

The heteromeric behavior we observe provides a plausible model for how higher-order heteromeric assemblies may have emerged during ancestral evolution. The most closely related example is the Family 2B class of two-component encapsulins, which shares the same HK97-fold structure as Family 1 QtEnc, but features an extensive insertion domain that is postulated to regulate the biosynthesis of secondary metabolites (*44*). Critically, our evolved heteromers mirror the assembly behavior of a representative Family 2B encapsulin recently characterized from *Streptomyces lydicus* (*45*). This encapsulin is encoded as two distinct copies in operon format, forming heteromeric complexes in which one of the two copies is assembly-incompetent. More generally, the capsids of adeno-associated viruses (AAVs) are also known to form stochastic assemblies of variable stoichiometry (*46*), which may influence native infectivity as well as transduction efficiency for gene therapy applications (*47*).

We anticipate that hybrid diversification arising from gene duplication represents a generally applicable concept for the directed evolution of homomeric proteins. This expectation is supported by other non-evolutionary protein engineering studies that demonstrate rescue of defective assemblies by WT co-expression (*48, 49*). While gene duplication holds significant potential for the optimization of viral delivery vectors, our strategy is perhaps most transformative for evolving non-viral multimeric proteins (*50*). Unlike the directed evolution of virus-like assemblies (*51, 52*), gene duplication expands access to diverse sequences without requiring encapsulation of encoding genetic information. Hence, our successful implementation of dual-compensatory gene duplication lays the foundation for addressing a wide variety of previously intractable biotechnological and biomedical challenges in the field of directed evolution.

## Acknowledgments

We thank S. Whitney for his insightful comments and discussion around this manuscript, as well as S. Pulsford and C. Jackson for their feedback. We acknowledge the core facilities from Sydney Analytical for infrastructure support and Sydney Microscopy and Microanalysis for microscopy support.

## Funding

Australian Research Council Discovery Project grant DP230101045 (YHL)

National Institutes of Health (R35GM133325) (TWG)

National Science Foundation (grant number: 2342136) (TWG)

## Author contributions

Conceptualization: RS, FL, YHL

Methodology: RS, FL, MPA, TWG, YHL

Investigation: RS, FL, TNS, AL, MPL, TWG

Visualization: RS, FL, AL, TWG, YHL

Funding acquisition: TWG, YHL

Project administration: RS, FL, TWG, YHL

Supervision: RS, TNS, YHL

Writing – original draft: RS, FL, TWG, YHL

Writing – review & editing: RS, TWG, YHL

## Data and materials availability

All data from this study are available from the corresponding authors upon request. Raw NGS reads from Illumina sequencing will be accessible on NCBI Sequence Read Archive at the point of publication. Source data for figures are available in supplementary tables data file. The rest of the data are available in the paper and supplementary materials. All analysed NGS datasets are available on https://github.com/LauGroup/EncDE/Selections and will be accessible at the point of publication.

## Code availability

All code generated for this study is available on https://github.com/LauGroup/EncDE and will be accessible at the point of publication.

