## Supplementary Materials for "Directed evolution of multimeric proteins is enabled by dual-compensatory gene duplication"

**This PDF file includes:**

Materials and Methods
Figs. S1 to S17
References

### Materials and Methods

#### Reagents

Commercially available reagents are listed with catalog numbers in Table S1. Oligonucleotides and plasmids used in this study are listed in Table S2 and S3 respectively.

#### Molecular cloning

Chemically competent *E. coli* DH5α cells were used for molecular cloning and plasmid propagation. DNA templates for cloning were synthesised from IDT (Integrated DNA Technologies). DNA amplification for cloning was done using Phusion™ polymerase (New England Biolabs; M0530). Plasmids used in this study were cloned by Gibson assembly (*1*) using NEBuilder HiFi DNA Assembly Master Mix (New England Biolabs; M5520AA) or purchased from Genscript unless otherwise specified. Gene sequences were verified by Sanger sequencing at the Australian Genome Research Facility. All details of plasmids, their descriptions and sequences are listed in Table S3.

#### *E. coli* growth assays

Growth assays were performed in electrocompetent *E. coli* BL21(DE3) cells under chloramphenicol (CAM) selection pressure as a measure of total chloramphenicol acetyltransferase (CAT) activity in cells. 50 ng of each plasmid was electroporated into 50 µL of cells, followed by recovery with 300 µL SOC (Super Optimal broth with Catabolite repression) media at 37 °C for 3 hours with shaking at 200 rpm. 100 µL of the recovery mixture was inoculated into 5 mL of LB (lysogeny broth) and incubated overnight for a further 16 hours. Transformed strains containing plasmids in pExplorer format only (pETDuet-1 backbone) were grown with ampicillin (100 µg/mL). Co-transformed strains containing plasmids in both pCompensator (pCDFDuet-1 backbone) and pExplorer format were grown with ampicillin (100 µg/mL) and spectinomycin (50 µg/mL). After overnight growth, the cultures were diluted with MilliQ water to an $OD_{600}$ of 0.5 to normalize the number of cells.

Spot assays on agar plates were performed by preparing 1:10 serial dilutions of the normalized cultures in MilliQ water. 3 µL of each serial dilution was spotted onto agar plates that were supplemented with IPTG (isopropyl β-D-1-thiogalactopyranoside, 0.1 mM) and CAM (0, 3, 5 or 10 µg/mL) to apply selection pressure.

Growth curves were measured by diluting the overnight cultures to an $OD_{600}$ of 0.1 with LB supplemented with IPTG (0.1 mM) and CAM (0, 5, 10 or 15 µg/mL), in a final volume of 100 µL. For strains with pCompensator included, spectinomycin (50 µg/mL) was also added. Cells were grown in 96-well clear flat bottom polystyrene microplates (Corning; 3370) at 37 °C with shaking at 200 rpm, measuring the $OD_{600}$ every 5 mins using a PHERAstar FSX multimodal plate reader (BMG Labtech). Growth curves were plotted using GraphPad Prism 10 software and presented as the mean of three technical replicates with standard deviation.

Plasmid library construction in pExplorer

The gene for QtEnc WT was first cloned into pETDuet-1 with CAT-CLP-ssrA in pExplorer format to be used as a template for further library construction.

To construct the Gen 1 library (Left/Right deletion), oligonucleotides were systematically designed with incremental binding regions towards either the left or right of the QtEnc WT gene, starting from a fixed point. The binding region of each oligonucleotide was preceded at the 5'-end by one randomized NNK codon and a Gibson overhang sequence (Fig. S1). To pair with these oligonucleotides, two additional oligonucleotides were designed to bind the AmpR gene in opposite directions. Given that PCR of each oligonucleotide pair generates fragments of incrementally smaller size, they were not pooled. Rather, each PCR was performed individually to avoid a nested PCR which would introduce library bias by favouring smaller fragments. Therefore, to create the 'Left' library, 14 oligonucleotides representing each deletion towards the left were paired with 1 AmpR oligonucleotide to generate 14 deletion-containing PCR products, covering the longer portion (~5000 bp, the variable fragments) of the final plasmid. The other shorter portion (~2000 bp, the constant fragment) was created by an oligonucleotide with compatible Gibson overhang on the QtEnc gene and another oligonucleotide with compatible overhang on the AmpR gene. The same process was applied to the right side with 15 oligonucleotides to create the 15 fragments for the 'Right' library. The oligonucleotides used to generate this library are listed in Table S2.

PCR was performed in a final reaction volume of 10 μL using Phusion™ reaction buffer, 0.2 mM of dNTPs, 0.5 μM of each primer, and 1 U of Phusion™ polymerase (New England Biolabs; M0530) and 4 ng of template DNA. Cycling conditions for the longer variable fragments were a denaturation cycle (98 °C, 30 s), followed by 35 cycles of denaturation (98 °C, 10 s), annealing (63 °C, 30 s), extension (72 °C, 3 min), and a final cycle of amplification (72 °C, 6 min). Cycling conditions for the shorter constant fragment were a denaturation cycle (98 °C, 30 s), followed by 35 cycles of denaturation (98 °C, 10 s), annealing (60 °C, 30 s), extension (72 °C, 1 min), and a final cycle of amplification (72 °C, 2 min). The amplicons were run on a 1% agarose gel (Sigma-Aldrich; A0169) with 1 kb Plus DNA ladder (New England Biolabs; N3200S) to verify their sizes. 2 μL of each of the variant fragments were pooled. 10 μL of the variant fragment pool was mixed with 10 μL of constant fragment and digested with 10 U of DpnI (New England Biolabs; R0176S), in 1X Cutsmart buffer (New England Biolabs; B7204S) at 37 °C for 2 hours to remove the template plasmid. The resulting digest was purified using an ISOLATE II PCR and Gel Kit (Bioline; BIO-52060) and eluted in 20 μL of MilliQ water. The fragments were ligated by mixing 20 μL of the purified fragment pool with 20 μL of 2X Gibson reaction mix (NEBuilder HiFi DNA Assembly Master Mix; M5520AA). The Gibson reaction mixture was split into 4 tubes to minimize bias and incubated at 50 °C for 4 hours to assemble all fragments. The assembled fragments were pooled and purified again using an ISOLATE II PCR and Gel Kit (Bioline; BIO-52060) and eluted in 15 μL of MilliQ water. To propagate the assembled plasmid library, 5 μL was electroporated into 100 μL of electrocompetent *E. coli* DH5α cells and recovered in 300 μL of SOC media for 2.5 hours at 37 °C and 200 rpm. To minimize bias, 2 aliquots of cells were electroporated per library. The total recovery was pooled and inoculated into 25 mL of LB supplemented with ampicillin (100 μg/mL) and grown overnight at 37 °C with shaking at 200 rpm for 16 hours. Cultures were harvested by centrifuging at 3000 g for 20 min, and plasmid libraries were isolated using ISOLATE II Plasmid Mini Kit (Bioline; BIO-52057) and eluted in 50 µL of MilliQ water. Both 'Left' and 'Right' deletion libraries were generated in this manner and mixed in a 1:1 ratio to create the Left/Right library.

To construct the Gen 2 library (Middle-Out deletion library), oligonucleotides were systematically designed with incremental binding regions towards the left or right of the QtEnc WT gene, starting from a fixed point. At the 5'-end of each oligonucleotide, an additional linker sequence was added, encoding a GSGGSG linker, which also act as Gibson overhangs (Fig. S2). Another oligonucleotide was designed binding the AmpR gene in the opposite direction of each set of deletion oligonucleotides. For the same reason as above, to avoid nested PCR bias favouring smaller fragments, individual PCR amplifications were performed. Each PCR would create one portion of the plasmid while introducing deletions in QtEnc gene and inserting the linker sequence. 14 individual reactions were performed to create 14 deletion fragments towards the left direction, creating the longer portion of the plasmid (~5000 bp). 15 individual reactions were performed to create deletion fragments towards the right direction, creating the shorter portion of the plasmid (~2000 bp). The oligonucleotides used to generate this library are listed in Table S2. All PCR fragments had the same linker sequence in the QtEnc gene, and overlapping AmpR sequence, serving as Gibson overhangs. Therefore, all fragments were pooled and a Gibson assembly reaction was performed, generating plasmids with deletions from the middle-out in both directions, bridged by a linker. All fragment amplification by PCR, digestion with DpnI, assembly by Gibson reaction, and plasmid library propagation, was performed as described above for the Gen 1 library.

To construct the Gen 3 library (Δ13 deletion library), oligonucleotides were designed to create a 19-residue deletion in the QtEnc gene (residues 190-208) while adding back six randomized NNK codons into the deleted region, as well as a compatible Gibson overhang (Fig. S3). The $(NNK)_6$ oligonucleotide pool was paired with an AmpR-binding oligonucleotide in the opposite direction. The oligonucleotides used to generate this library are listed in Table S2. PCR amplification created one portion of the plasmid. PCR was performed as described previously, but instead using Q5® High-Fidelity 2X Master Mix (New England Biolabs; M0492S) with an initial denaturation cycle (98 °C, 30 s), followed by 35 cycles of denaturation (98 °C, 10 s), annealing (64 °C, 30 s), extension (72 °C, 1 min), and a final cycle of amplification (72 °C, 2 min). The other portion of the plasmid was created as before using oligonucleotides binding to the QtEnc gene and the AmpR gene in the other direction with compatible Gibson overhangs. PCR was performed with an initial denaturation cycle (98 °C, 30 s) followed by 35 cycles of denaturation (98 °C, 10 s), annealing (64 °C, 30 s), extension (72 °C, 3 min), and a final cycle of amplification (72 °C, 6 min). 10 µL of each amplified fragment was digested with 5 U of DpnI in 1X Cutsmart buffer at 37 °C for 2 hours to remove the template plasmid, then purified as described above. Gibson assembly was performed by mixing 7.5 µL of each of the two fragments with 15 µL of 2X Gibson reaction mix (NEBuilder HiFi DNA Assembly Master Mix; M5520AA). The reaction was split into 3 tubes to minimize bias and incubated at 50 °C for 4 hours to assemble all fragments into plasmids. The assembled plasmids were pooled, purified and propagated to create the final library as described above.

To construct the Gen 4 library (Mixed library), the Gen 1 (Left/Right deletion libraries) and Gen 2 (Middle-Out library) were first shuffled by StEP recombination (*2*) (Fig. S4). Oligonucleotides were designed to amplify the variable segment of the QtEnc gene in each library as listed in Table

S2 to create the shuffled insert. PCR was performed in a final reaction volume of 10 µL using 0.5 µM of each primer, One*Taq*® 2X Master Mix (New England Biolabs; M0482S) and 25 ng of template DNA. Cycling conditions involved an initial denaturation cycle (98 °C, 30 s) followed by 99 cycles of denaturation (98 °C, 10 s), annealing (55 °C, 5 s), extension (50 °C, 5 s). The shuffled amplicon was digested with 5 U of DpnI in 1X Cutsmart buffer at 37 °C for 2 hours to remove the template. 1 µL of this amplicon was used as a template for a second bump-up PCR to amplify the shuffled insert. This PCR was performed in a final reaction volume of 10 µL using 0.5 µM of the same primers, with Q5® High-Fidelity 2X Master Mix with an initial denaturation cycle (98 °C, 30 s), followed by 35 cycles of denaturation (98 °C, 10 s), annealing (68 °C, 30 s), extension (72 °C, 30 s), and a final cycle of amplification (72 °C, 1 min). The backbone for this library was created by designing oligonucleotides flanking the variable region of QtEnc gene that are outward facing. PCR amplification therefore results in a long fragment (~7000 bp) including the plasmid backbone. This PCR was performed in a final reaction volume of 10 µL using Q5® High-Fidelity 2X Master Mix, 0.5 µM of each primer, and 1 ng of template DNA. Cycling conditions involved an initial denaturation cycle (98 °C, 30 s), followed by 35 cycles of denaturation (98 °C, 10 s), annealing (68 °C, 30 s), extension (72 °C, 3 min 30 s), and a final cycle of amplification (72 °C, 7 min). 10 µL of this backbone fragment was DpnI digested and purified as described above. Gibson assembly was performed by mixing the 7.5 µL of the insert with 7.5 µL of the backbone fragment and 15 µL of 2X Gibson reaction mix as described above. The assembled plasmids were purified and propagated to create the shuffled plasmid library as described above. This shuffled library, the Left/Right library (Gen 1), Middle-Out library (Gen 2), and Δ13 deletion library (Gen 3) were mixed together in a 1:1:1:1 ratio to create the final 'Mixed' library.

Library selection assays in *E. coli*

Selection experiments were performed in electrocompetent *E. coli* BL21(DE3) cells with plasmid libraries in pExplorer format (pETDuet-1 backbone) with chloramphenicol (CAM) selection pressure. 100 ng of pExplorer plasmid library was transformed by electroporation into 100 µL of cells. Selections were also performed on co-transformed strains containing pExplorer and pCompensator with CAM to apply selection pressure and spectinomycin to maintain the pCompensator (pCDFDuet-1 backbone) plasmid. Co-transformations were performed by mixing 100 ng of pExplorer plasmid library and 100 ng of pCompensator plasmid and electroporated into 100 µL of cells. Post electroporation, cells were recovered with 1 mL SOC media at 37 °C for 3 hours with shaking at 200 rpm. 150 µL of the recovery mixture was inoculated into 5 mL of LB supplemented with IPTG (0.1 mM), CAM (0, 5, 10, 15 or 20 µg/mL) and incubated at 37 °C for 20 hours with shaking at 200 rpm. For co-transformations with pCompensator, spectinomycin (50 µg/mL) was also included. Each selection was performed as experimental duplicates. If growth was observed, the cultures were harvested at stationary growth phase ($OD_{600}$ >1) by centrifugation at 3000 g for 20 min, and plasmids were isolated using ISOLATE II Plasmid Mini Kit (Bioline; BIO-52057) (Fig. 2B).

Next-Generation Sequencing (NGS) of libraries before and after selection

To determine the distribution of deletions in the variable region of the QtEnc gene, each of the plasmid libraries was sequenced before and after selection pressure was applied. Plasmid DNA libraries were isolated as described above. A standard Illumina pipeline of two-step PCR was performed for short-read NGS. The first PCR reaction was performed using oligonucleotides flanking the variable segment of the QtEnc gene with adapter sequences on the ends compatible with indices from Nextera XT Index kit V2 (Illumina). The second PCR was performed with indexing oligonucleotides from Nextera XT Index kit V2 (Illumina) to attach the barcodes and Illumina P5 and P7 sequences. The first PCR was performed with 25 ng of template DNA, in a final reaction volume of 10 µL using 0.5 µM of the 'NGS primers' (Table S2), using Q5® High-Fidelity 2X Master Mix with an initial denaturation cycle (98 °C, 30 s) followed by 15 cycles of denaturation (98 °C, 10 s), annealing (68 °C, 30 s), extension (72 °C, 30 s), and a final cycle of amplification (72 °C, 1 min). 1 µL of the first PCR reaction was used as template DNA to setup the second PCR with 'index primers', with the cycling conditions involving an initial denaturation cycle (98 °C, 30 s), followed by 15 cycles of denaturation (98 °C, 10 s), annealing (59 °C, 30 s), extension (72 °C, 30 s), and a final cycle of amplification (72 °C, 1 min). Each PCR was performed in duplicate and pooled together after indexing and purified using an ISOLATE II PCR and Gel Kit, eluting in 15 µL of MilliQ water. Samples were prepared for sequencing on an Illumina iSeq-100 instrument (single-read) with read length of 151 bp. The list of adapters and oligonucleotides used to perform the two step PCR are in Table S2.

Selection enrichment and fold-change analysis

Demultiplexing and adapter trimming was performed using Illumina iSeq local run manager software to create FASTQ reads. FASTQ files were processed and analysed using a series of custom Python scripts available on https://github.com/LauGroup/EncDE with detailed instructions and all datasets. The following Python packages: pandas (*3*), NumPy (*4*), SciPy (*5*), Matplotlib (*6*), Logomaker (*7*), openpyxl (*8*) and software: Clustal Omega v1.2.4 (*9*) were used in the analysis pipeline. Reads were first filtered to contain the correct flanking sequences surrounding the variable region of interest. Each DNA read was translated to amino acids and clustered by unique sequences. Number of unique sequences detected in each selection with and without pCompensator was compared (Fig. 2C). Resulting unique sequences were aligned using Clustal Omega multiple sequence alignment software package (*9*) and number of deletions was counted in each sequence compared to QtEnc WT. Sequence logo representation of the library was generated using Logomaker package (*7*). The read count for each unique sequence was quantified. For each sample, % frequency was calculated for each sequence as (read count / total read count) x 100 and converted to $log_2$ as appropriate. Libraries before selection were represented as $log_2$ frequency in experimental duplicates to assess library bias. In addition, the distribution of deletions in each sample was visualized by coloring each sequence according to number of deletions compared to WT. The library after selection was sequenced and processed similarly. To calculate fold-change of each sequence after selection compared to before selection, we applied the standard practice of adding a pseudo-count of 1 to frequency and converting into reads per million (RPM) instead of percentage (*10*, *11*). Only sequences with raw read counts above 20 were considered to determine top performing sequences. Therefore, RPM for each sequence was calculated as (read count / total read count) x 10^6 and fold-change was calculated as (RPM after selection +1) / (RPM before selection +1) and converted to $log_2$ as appropriate.

When selection was performed with WT pCompensator, libraries after selection had approximately half of the total reads (~50% frequency) corresponding to the WT sequence as expected. However, WT reads were also present in sequencing results from libraries before selection, as well as after selection without WT pCompensator. These artefact WT reads arose from residual template DNA, only reflecting the variable degree of incomplete DpnI digestion. Thus, % frequency and fold-change calculations on WT reads in these instances were disregarded. Only non-WT sequences were considered as top performers and these were not compared against WT.

Frequency values for libraries before selection and fold-change values for all libraries after selection are available on https://github.com/LauGroup/EncDE/Selections.

Expression and purification of encapsulins assembled in *E. coli*

Proteins were expressed and purified as previously reported (*12*). Sequences of proteins purified are listed in Table S4. Chemically competent *E. coli* BL21(DE3) cells were transformed with plasmids and inoculated into 10 mL cultures of LB supplemented with ampicillin (100 µg/mL). After overnight incubation at 37 °C with shaking at 200 rpm, the cultures were used to inoculate 400 mL cultures of LB supplemented with ampicillin (100 µg/mL), grown at 37 °C with shaking at 200 rpm. When the $OD_{600}$ reached 0.6-0.8, protein expression was induced with IPTG (0.1 mM) and cultures were grown overnight at 18 °C. Cells were then harvested at 3214 g for 20 min (5810R centrifuge with S-4-104 rotor, Eppendorf) and stored at -20 °C ready for purification.

Proteins were purified by ammonium sulfate precipitation and multiple size exclusion steps. Cell pellets were resuspended in lysis buffer (50 mM Tris pH 8.0, 200 mM NaCl, 10 µg/mL DNase I (Sigma-Aldrich; AMPD1-1KT)), 100 µg/mL lysozyme (Sigma-Aldrich; 12650-88-3) and 1X protease inhibitor (cOmplete EDTA-free Protease Inhibtor Cocktail, Roche; 11836170001). Pellets were thawed on ice for 30 min and lysed by sonication using Sonopuls HD 4050 with TS-106 probe (Bandelin) for 11 min at 55% amplitude with a pulse time of 8 s on and 10 s off. Cell lysate was clarified by centrifugation at 17000 g for 40 min and the supernatant was separated. Solid ammonium sulfate (Sigma-Aldrich; A4418) was added to the supernatant to 20% (w/v) saturation and rocked at 4 °C for 15 min then centrifuged at 10000 g for 15 min and the supernatant was separated again. Solid ammonium sulfate was added to this supernatant again to a total of 50% (w/v) saturation with rocking at 4 °C for 15 min then centrifuged at 10000 g for 15 min. The supernatant was discarded and the pellet containing the proteins was kept and resuspended in 5 mL of size-exclusion chromatography (SEC) buffer (50 mM Tris pH 8.0, 200 mM NaCl), then dialysed using Snakeskin Dialysis Tubing (3.5 kDa MWCO, ThermoFisher Scientific; 88242) against 1 L of SEC buffer at 4 °C for 2-4 h followed by changing the buffer to another 1 L of fresh SEC buffer and left dialysing overnight at 4 °C. Post dialysis, the sample was recovered and subjected to anion-exchange chromatography to remove nucleic acid contaminants on a HiPrep Q XL 16/10 column at a flow rate of 5 mL/min in SEC buffer. The flowthrough containing protein was collected while the nucleic acid contaminants were retained on the column and discarded. The protein was further purified through two rounds of size-exclusion chromatography in SEC buffer. The first SEC was performed using a HiPrep 16/60 Sephacryl S-500 HR column and the fractions containing the protein was collected and concentrated using Amicon Ultra-15 filters 100 kDa MWCO (Merck; UFC9100). The concentrated sample was applied to Superose 6 Increase 10/300 GL column to perform the second SEC. The fractions containing purified protein were pooled, quantified via measuring absorbance at 280 nm and stored at 4 °C until used. The sample was analysed by SDS-PAGE (sodium dodecyl sulfate polyacrylamide gel electrophoresis) using Any kD Mini-PROTEAN TGX Stain-Free gels (Bio-Rad; 4568123, 4568126) with Tris-glycine-SDS running buffer (Bio-Rad; 1610772). For size comparison, Unstained Protein Standard, Broad Range 10-200 kDa (New England Biolabs; P7717S) was used.

Expression and purification of encapsulin fusions for *in vitro* assembly

Encapsulin fusion constructs that result in pre-assembly monomers are described in Table S3. Sequences of purified fusion proteins and cleaved proteins are listed in Table S4. The fusion proteins were expressed in *E. coli* using the same method as described above for *E. coli* assembled encapsulins. The proteins were purified by lysing the pellets by sonication as described above but in lysis buffer (50 mM Tris pH 8.0, 200 mM NaCl, 1 mM tris(2-carboxyethyl)phosphine (TCEP) (Merck; 646547), 10 µg/mL DNase I, 100 µg/mL lysozyme and 1X protease inhibitor). The clarified lysate was subjected to Ni-NTA affinity chromatography by applying to 1 mL of Ni-NTA resin (ThermoFisher Scientific; 88221) equilibrated in Ni-NTA buffer (50 mM Tris pH 8.0, 200 mM NaCl, 1 mM TCEP). The mixture was added to a gravity flow column and washed with 3 x 2 mL wash buffer (50 mM Tris pH 8.0, 200 mM NaCl, 1 mM TCEP, 20 mM imidazole) and eluted with 4 x 1 mL elution buffer (50 mM Tris pH 8.0, 200 mM NaCl, 1 mM TCEP, 500 mM imidazole). The eluted samples were pooled, analysed by SDS-PAGE and quantified as described above. The purified proteins were then buffer exchanged into storage buffer (50 mM Tris pH 8.0, 200 mM NaCl, 1 mM TCEP) by concentrating using Amicon Ultra-15 centrifugal filters 10 kDa MWCO ( Merck; UFC9010) up to 100 µM and snap frozen to -80 °C for storage until use.

*In vitro* assembly of homomeric and heteromeric hybrid encapsulins

The encapsulin fusion pre-assembly monomers were *in vitro* assembled into homomeric protein cages, triggered by cleavage of the fusion partner with TEV protease as previously reported (*12*).

Small scale cleavage reactions were performed on 60 µL scale by incubating 20 µM encapsulin fusions and 20 µM TEV protease in SEC buffer (50 mM Tris pH 8.0, 200 mM NaCl, 1 mM TCEP) overnight at room temperature or 25 °C. This sample was analysed on SDS-PAGE to verify if cleavage had occurred, while 40 µL was subjected to analytical SEC using a Bio SEC-5 2000 Å column to verify if the elution profile was consistent with a large protein cage being formed. Fractions were collected and concentrated using Amicon Ultra-0.5 centrifugal filters 100 kDa MWCO (Merck; UFC5100) and analysed by SDS-PAGE to verify the correct size.

For cryo-EM structure determination, the reaction was performed similarly but scaled up to 400 µL. After overnight cleavage, the reaction was directly subjected to size-exclusion chromatography on a Superose 6 Increase 10/300 GL column to purify the assembled cages. Heteromeric hybrid encapsulins were *in vitro* assembled similarly by incubating 20 µM assembly-competent monomers with a range of concentrations (5, 20, 40, 80 µM) of assembly-incompetent monomers and 20 µM TEV protease. The reactions were analysed and purified in the same manner.

Analytical size-exclusion chromatography (SEC)

Analytical SEC was performed using a Nexera Bio HPLC (Shimadzu) system with a Bio SEC-5 2000 Å pore size column (7.8 × 300 mm, Agilent) with a Bio SEC-5 2000 Å guard column (7.8 × 50 mm, Agilent). The mobile phase was SEC buffer (50 mM Tris pH 8.0, 200 mM NaCl, 1 mM TCEP) at a flow rate of 1 mL/min for 30 min, injecting 40 µL samples from *in vitro* assembly cleavage reactions.

Dynamic light scattering (DLS)

Samples of encapsulin assemblies were loaded onto Prometheus Panta (NanoTemper) at 0.5 mg/mL in SEC buffer (50 mM Tris pH 8.0, 200 mM NaCl, 1 mM TCEP) according to manufacturer instructions. Size Analysis function was used to perform isothermal DLS scans (10 acquisitions, 5 s each, 100 % Laser intensity at 25 °C).

Cryo-electron microscopy (cryo-EM)

*Sample preparation*

Purified samples of Glass9 and Letter11 were concentrated to 3.7 and 4.5 mg/mL respectively, in 200 mM NaCl, 25 mM Tris pH 8.0, 1 mM TCEP. 3.5 µL of protein samples were applied to freshly glow discharged Quantifoil R2/1 Cu 200 mesh grids and prepared by plunge freezing in liquid ethane using an FEI Vitrobot Mark IV (100% humidity, 22 °C, blot force 5, blot time 2 seconds). The grids were immediately clipped and stored in liquid nitrogen until data collection.

*Data collection*

Cryo-EM movies were collected using a ThermoFisher Scientific Titan Krios G3 cryo-electron microscope operating at 300 kV equipped with a Gatan K3 direct electron detector with a BioQuantum imaging filter. SerialEM (*13*) was used to select targets and acquire movies. See Table S5 for additional data collection statistics for each dataset.

*Data processing*

Glass9: CryoSPARC 4.7.1 (*14*) was used to process the dataset. 1,820 movies were imported, motion corrected by patch motion correction, and the CTF fit was estimated using patch CTF estimation in CryoSPARC *Live*. Blob picker was utilized to select 320 initial particles that were used to create templates for template-based particle picking. Template picker was used to select 68,389 particles which were then extracted using a box size of 684 pixels. The particles were then sorted by two rounds of 2D classification, resulting in 61,300 selected particles. These particles were then used to generate an initial volume by ab-initio reconstruction using three classes and I symmetry, resulting in a majority class containing 61,104 particles. The particles were then downsampled to a box size of 480 pixels and used for homogeneous refinement using the ab-initio map with I symmetry imposed, per-particle defocus optimization, per-group CTF parameterization, spherical aberration fitting enabled, tetrafoil fitting enabled, anisotropic magnification fitting enabled, and Ewald sphere correction enabled using a positive curvature sign, resulting in a map with 2.40 Å global resolution.

Letter11: CryoSPARC 4.7.1 (*14*) was used to process the dataset. 1,565 movies were imported, motion corrected by patch motion correction, and the CTF fit was estimated using patch CTF estimation in CryoSPARC *Live*. 237 particles were manually selected and used to make templates for particle picking. Template picker was used to select 68,389 particles which were then extracted using a box size of 684 pixels. The particles were then sorted by two rounds of 2D classification resulting in 68,033 selected particles, with 44,984 particles of T=4 icosahedral symmetry and 18,343 particles of T=3 icosahedral symmetry which were processed separately in subsequent steps. T=3 and T=4 particles were separately used to generate initial volumes for either T=3 and T=4 assemblies by ab-initio reconstruction using three classes and I symmetry, resulting in majority classes containing 44,749 particles (T=4) and 18,327 particles (T=3). The particles were then downsampled to a box size of 480 pixels and used for homogeneous refinement using the ab-initio maps with I symmetry imposed, per-particle defocus optimization, per-group CTF parameterization, spherical aberration fitting enabled, tetrafoil fitting enabled, anisotropic magnification fitting enabled, and Ewald sphere correction enabled using a negative curvature sign, resulting in a maps with 2.40 Å global resolution (T=4) and 2.42 Å global resolution (T=3).

*Model building*

For building the Glass9 and Letter11 models, starting models of individual protomers were generated using AlphaFold3 (*15*) and were placed manually into the respective maps using ChimeraX v.1.8 (*16*), followed by using the fit-in-map command. This step was repeated for the remaining three protomers in the T=4 maps and two protomers in the T=3 map, resulting in complete asymmetric units containing four protomers (Glass9 and Letter11 T=4) or three protomers (Letter11 T=3). The models were then manually refined against the density maps using Coot v0.9.8.1 (*17*). Phenix v 1.20.1-4487-000 (*18*) was then used to further refine the models by real-space refinement with three macrocycles, minimization_global enabled, local_grid_search enabled, and adp refinement enabled. NCS operators were identified from the map using map_symmetry and applied to the model using apply_ncs to generate NCS-expanded partial shells. The NCS-expanded partial shells were then refined again using real-space refinement with three macrocycles, minimization_global enabled, local_grid_search enabled, adp refinement enabled, and NCS constraints enabled. The BIOMT operators were identified using the find_ncs command and manually placed into the header of the .pdb files containing a single ASU of the NCS-refined models.

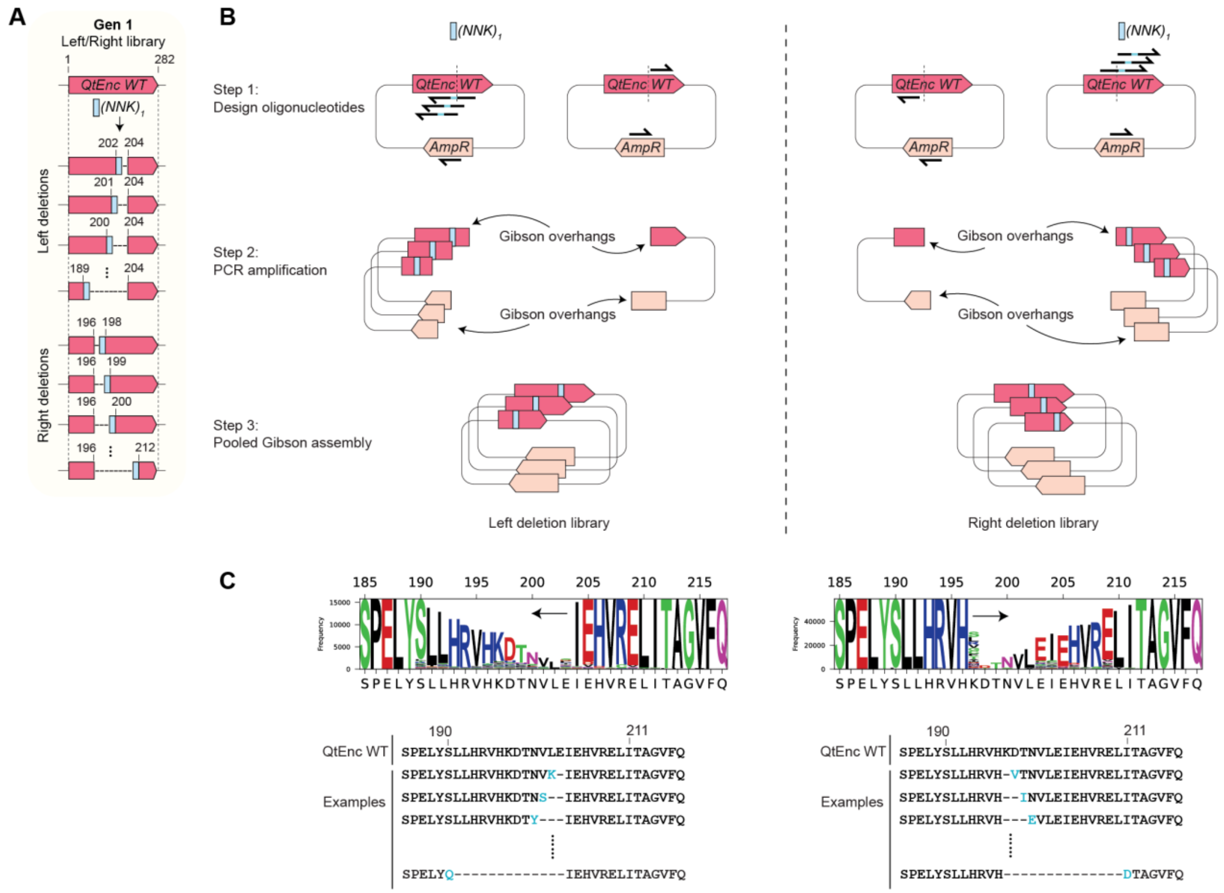


**Fig. S1. Gen 1 Left/Right deletion library construction.** (**A**) Schematic showing representative distribution of deletions in QtEnc WT gene. Note that numbering refers to the corresponding amino acid residues encoded by the gene. (**B**) Overview of cloning procedure for Left/Right deletion libraries. Oligos were systematically designed with incremental binding regions towards the left or right of the gene, preceded at the 5'-end by one randomized NNK codon and a Gibson overhang sequence. Each of the two deletion oligo sets (Left or Right) was paired with an AmpR-binding oligo in the opposite direction. Each PCR amplification created one portion of the plasmid while introducing deletions in the QtEnc gene. The other portion of the plasmid was amplified with compatible Gibson overhangs. Pooled Gibson assembly was performed on each pair of matching PCR fragments to generate each of the two plasmid libraries (Left or Right), which were then mixed in a 1:1 ratio to create the final Left/Right library. (**C**) Sequence logo of the Left and Right libraries shows successful library construction. Example sequences below show the incremental deletion window (-), with the variable residue at the end of each deletion shown in blue.

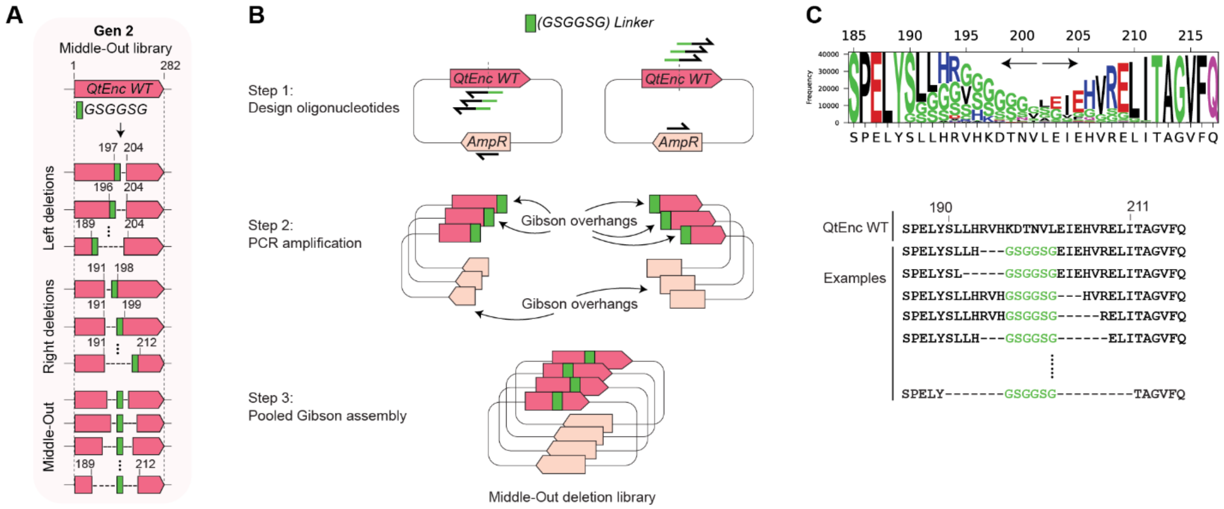


**Fig. S2. Gen 2 Middle-Out deletion library construction.** (**A**) Schematic showing representative distribution of deletions in QtEnc WT gene with an additional linker sequence encoding for a GSGGSG linker. Note that numbering refers to the corresponding amino acid residues encoded by the gene. (**B**) Overview of cloning procedure for Middle-Out deletion libraries. Oligos were systematically designed with incremental binding regions towards the left or right of the gene, with the linker sequence at the 5'-end that also acts as a Gibson overhang sequence. Each deletion oligo set was paired with an AmpR-binding oligo in opposite direction. Each PCR amplification created one portion the plasmid while introducing deletions in the QtEnc gene and insertion of the linker. Pooled Gibson assembly was performed to generate the Middle-Out library. (**C**) Sequence logo of the Middle-Out library shows successful library construction. Example sequences below show the range of deletions (-), with the GSGGSG linker shown in green.

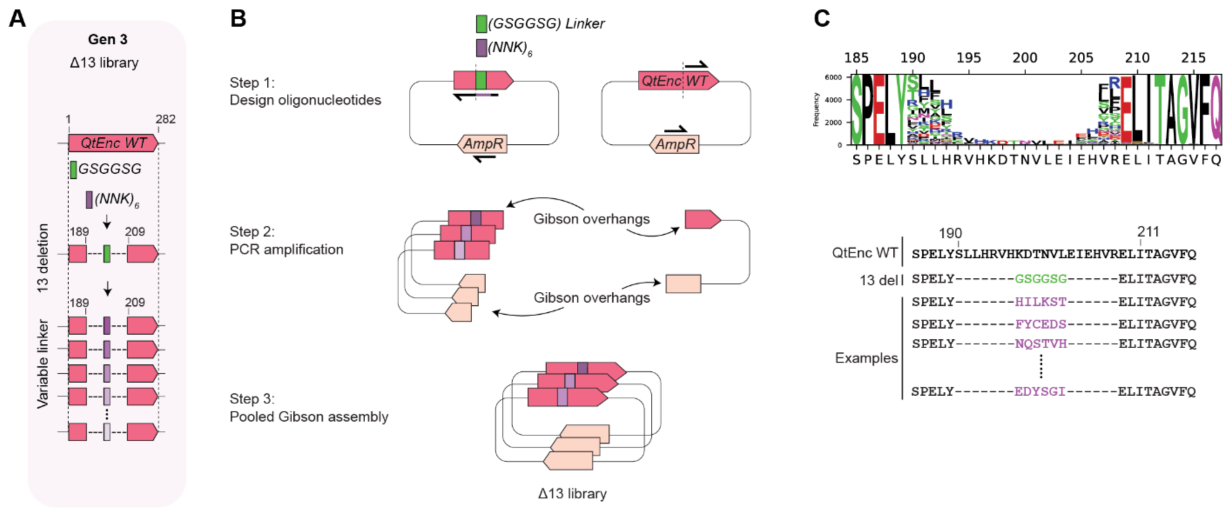


**Fig. S3. Gen 3 Δ13 deletion library construction.** (**A**) Schematic showing replacement of the GSGGSG linker with a variable linker on a QtEnc variant with 13 residues deleted. Note that numbering refers to the corresponding amino acid residues encoded by the gene. (**B**) Overview of cloning procedure for Δ13 deletion library. Oligos were designed with six randomized NNK codons to replace the GSGGSG linker sequence, as well as a compatible Gibson overhang. The $(NNK)_6$ oligo pool was paired with an AmpR-binding oligo in the opposite direction. PCR amplification created half of the plasmid while introducing the variable linker. The other half of the plasmid was amplified with compatible Gibson overhangs. Pooled Gibson assembly was performed to generate the Δ13 library. (**C**) Sequence logo of Δ13 library shows variable residues as intended. Example sequences from the library show all sequences have 13 deletions (-) and a variable six-residue linker shown in purple.

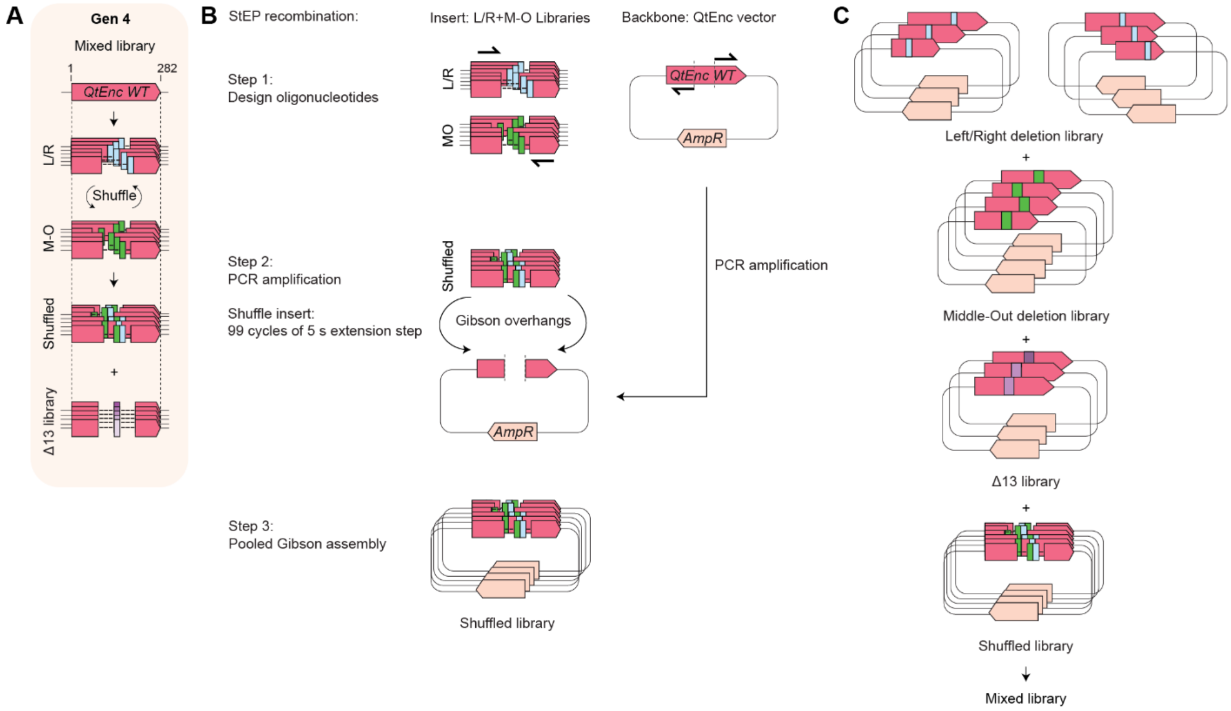


**Fig. S4. Gen 4 Mixed library construction.** (**A**) Schematic showing representative libraries contained in the Gen 4 mixture. (**B**) Overview of cloning procedure for gene shuffling Left/Right (L/R) and Middle-Out (M-O) deletion libraries by StEP recombination. Oligos were designed to bind the regions flanking the variable segment of the QtEnc gene in each library. The shuffled insert library was created by PCR amplification, with short 5 s extension steps over 99 cycles to recombine the two libraries. The plasmid backbone was generated by PCR amplification using oligos that bind to the same flanking regions of the QtEnc gene, generating Gibson overhangs matching those of the insert library. Pooled Gibson assembly was performed to generate the shuffled plasmid library. (**C**) All four libraries were mixed in a 1:1:1:1 ratio to create the final Mixed library.

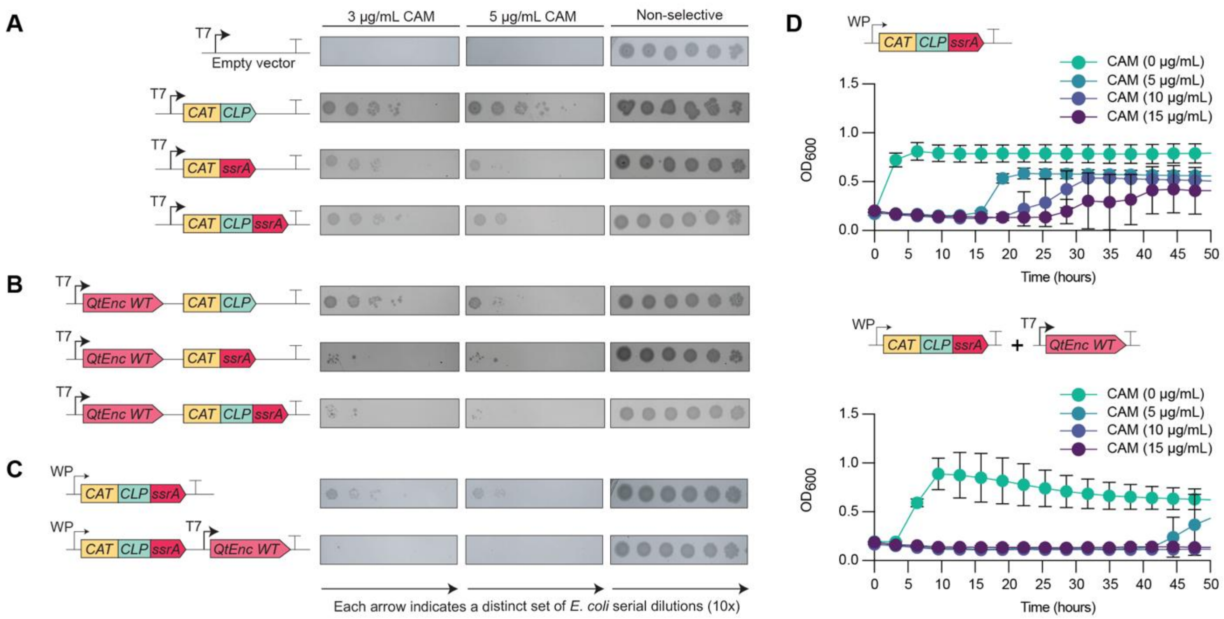


**Fig. S5. CAT promoter optimization and the inclusion of QtEnc WT achieves minimal growth under antibiotic selection.** (**A**) *E. coli* cells expressing different components of the selection circuit under strong T7 promoter spotted on selective plates (3 or 5 µg/mL CAM, 0.1 mM IPTG). While the ssrA tag does reduce survival, background growth remains too high. (**B**) Strains co-expressing QtEnc WT show lower background survival. The reduction in overall CAT activity is likely due to a combination of increased metabolic burden and CAT encapsulation. (**C**) Changing to a weak promoter (WP) for CAT achieves minimal background survival when QtEnc WT is included. (**D**) Growth curves for *E. coli* strains in liquid culture with different levels of selective pressure (0-15 µg/mL CAM, 0.1 mM IPTG) show that the high background of growth when expressing CAT-CLP-ssrA is supressed when QtEnc WT is included.

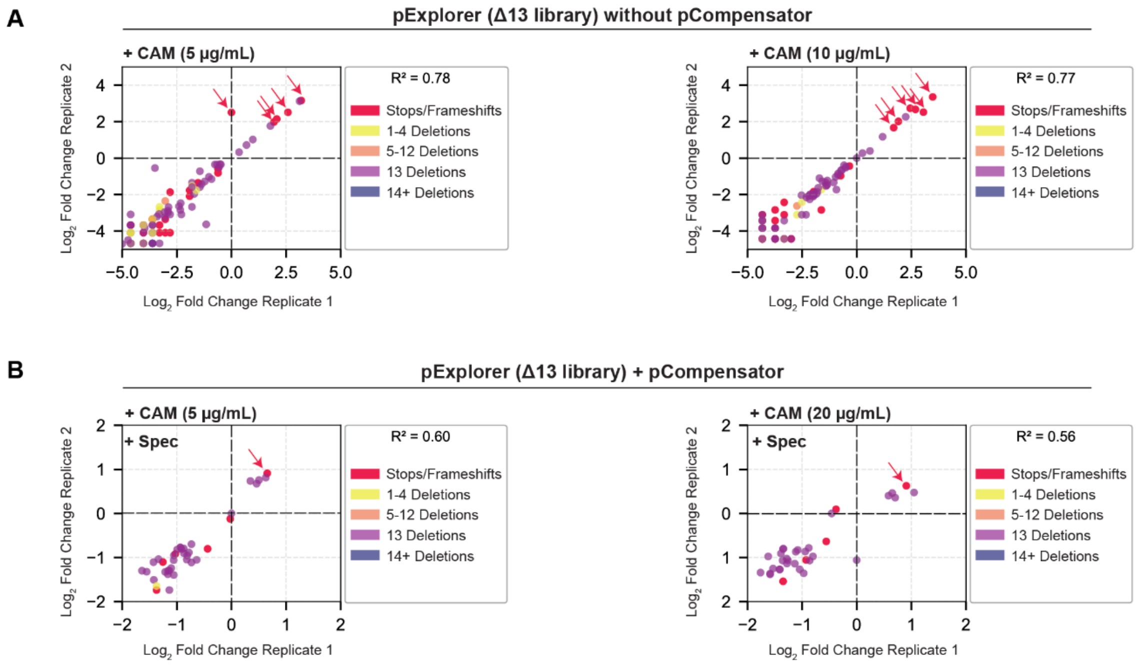


**Fig. S6. Using pCompensator during Δ13 library selection lowers the rate of false-positive stop codons and frameshifts.** $Log_2$ plots of fold change over input library are shown for experimental replicates with selection either (**A**) without pCompensator (5-10 µg/mL CAM, 0.1 mM IPTG) or (**B**) with pCompensator (5-20 µg/mL CAM, 50 µg/mL Spec, 0.1 mM IPTG). Without pCompensator, no growth was observed above 10 µg/mL CAM. Each encapsulin variant is represented as a single dot, colored according to number of deletions in each variant sequence, while QtEnc WT is shown in black. Enriched variants (top-right quadrant) with stop codons or frameshifts are colored in red and emphasised with a red arrow. These enriched false positives are more frequent without pCompensator (11 out of 19, i.e. 58%) than with pCompensator (2 out of 10, i.e. 20%).

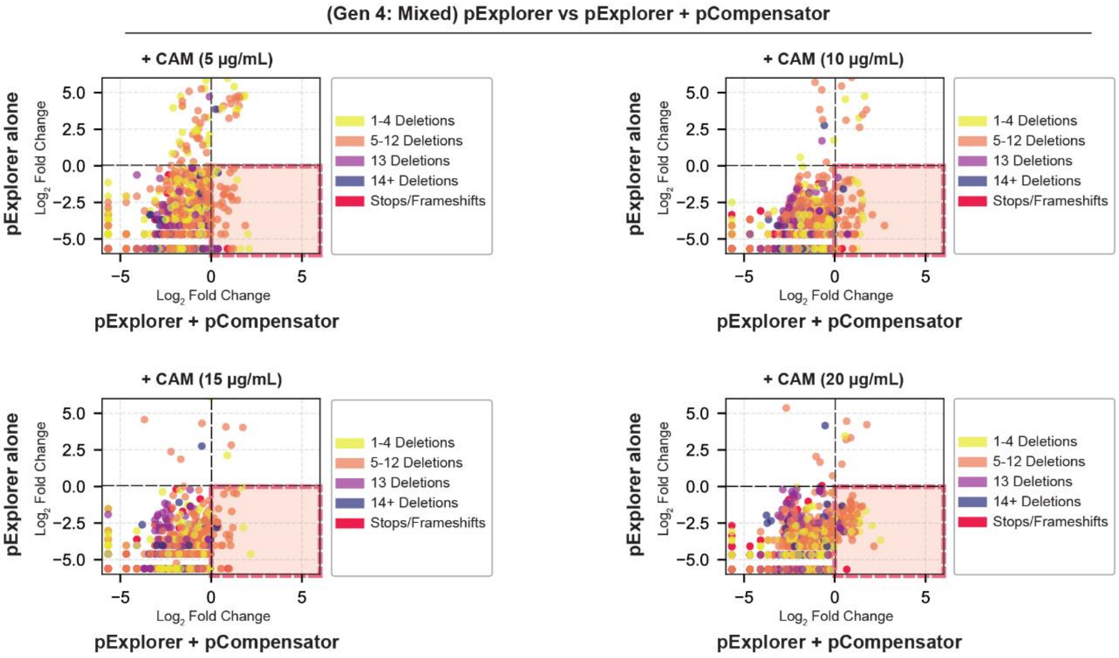


**Fig. S7. Selection of pExplorer (Gen 4: Mixed) with pCompensator yields unique sequences.** $Log_2$ plots of fold change over input library are shown for selection with pCompensator (5-20 µg/mL CAM, 50 µg/mL Spec, 0.1 mM IPTG) and without pCompensator (5-20 µg/mL CAM, 0.1 mM IPTG). Each encapsulin variant is represented as a single dot whose x-coordinate reflects fold change over input library after selection when pCompensator is included, and y-coordinate reflects fold change when pCompensator is absent. Sequences in the lower right quadrant (highlighted in pink) were only enriched with pCompensator. Dot points are colored according to number of deletions in particular variant, while QtEnc WT is shown in black. Variants selected for further study are outlined in black.

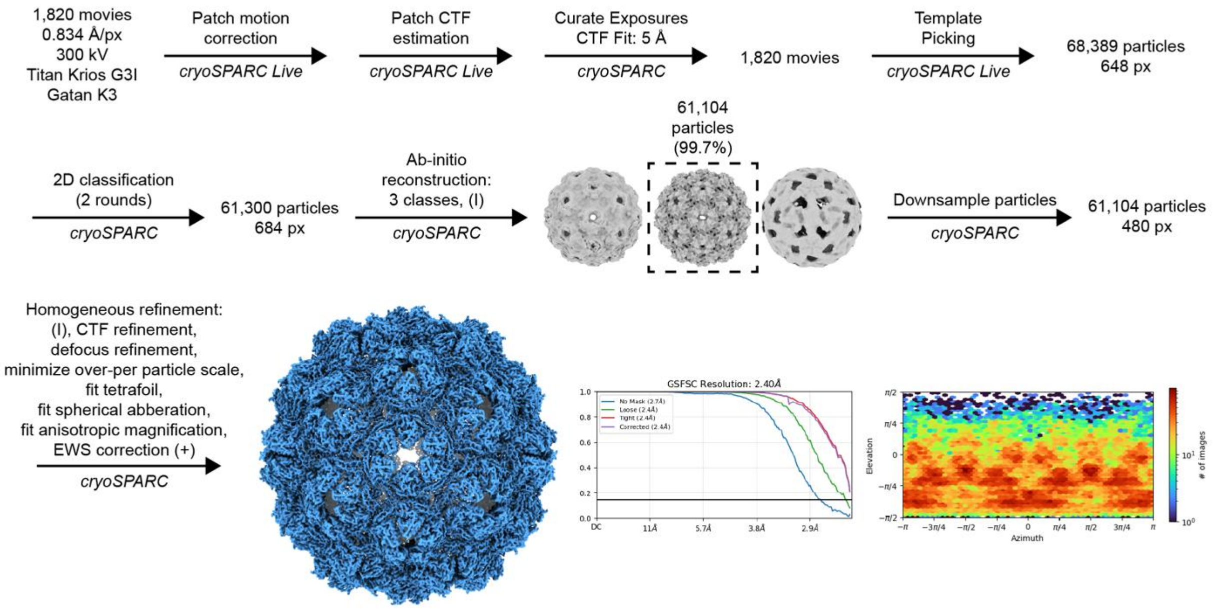


**Fig. S8. Cryo-EM data processing workflow for Glass9.** Representative micrographs and 2D class averages are shown in the main text (Fig. 4B).

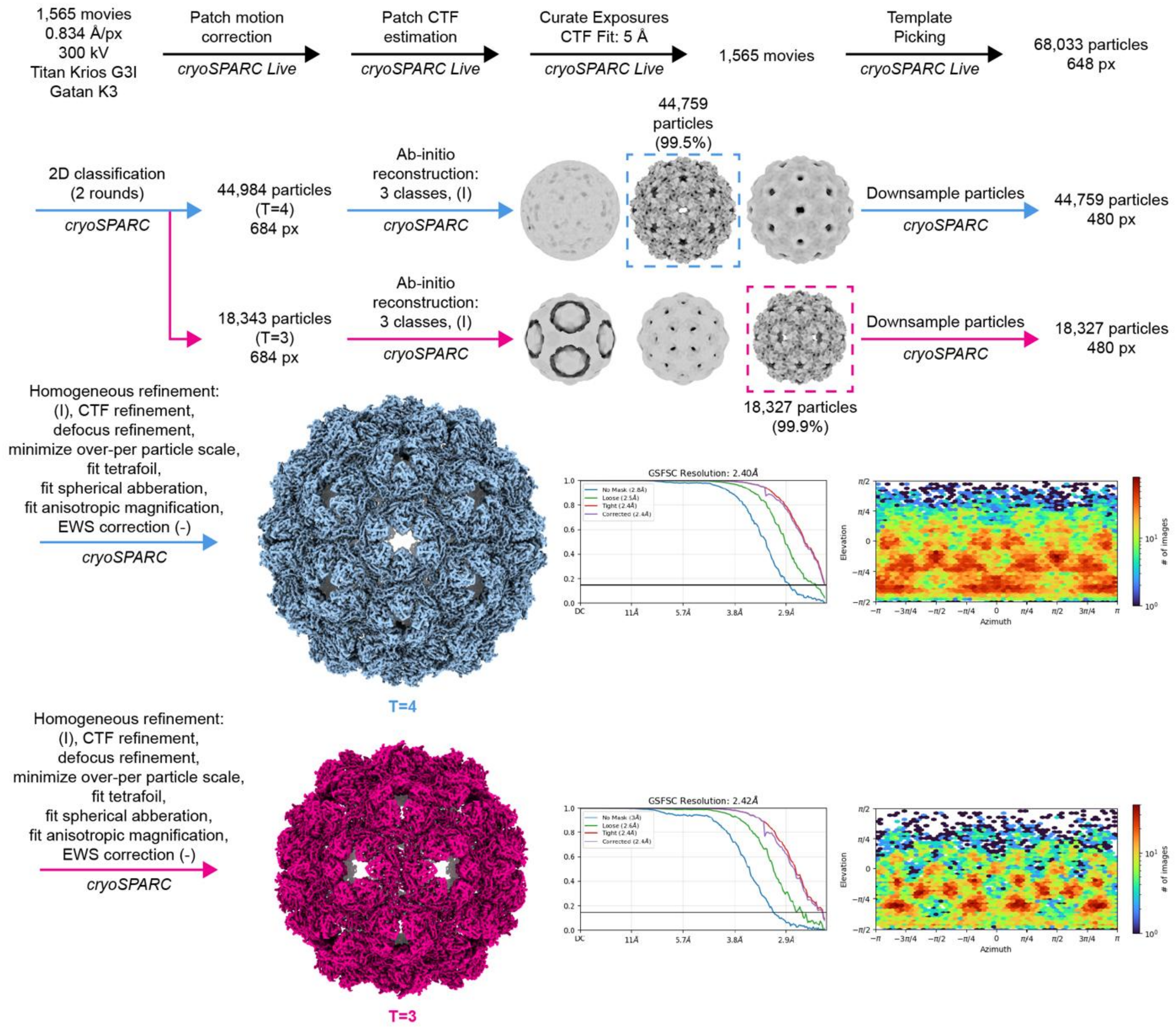


**Fig. S9. Cryo-EM data processing workflow for Letter11.** Representative micrographs and 2D class averages are shown in the main text (Fig. 4B).

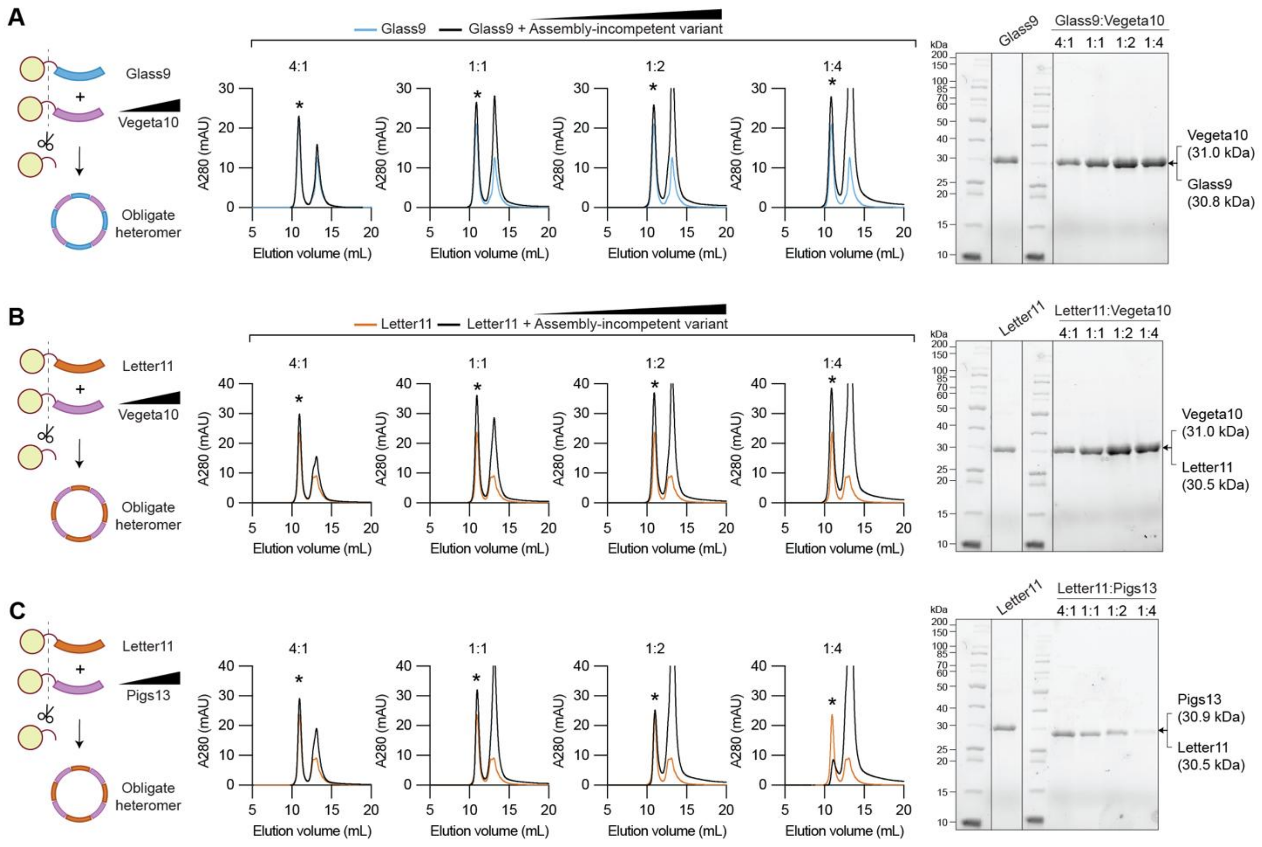


**Fig. S10. Assembly-competent variants can rescue assembly-incompetent variants to create variant-variant hybrid cages.** (**A-C**) Titration of pre-assembly monomers of assembly-competent variants (Glass9, Letter11; 20 µM) titrated with assembly-incompetent variants (Vegeta10, Pigs13; 5, 20, 40, 80 µM), followed by *in vitro* assembly. Size-exclusion chromatography on a Bio SEC-5 2000 Å HPLC column confirm the expected elution peak for hybrid cages (marked with asterisks). SDS-PAGE analysis of the expected elution peak for encapsulin assembly shows increased band intensity as both monomers have similar molecular weight.

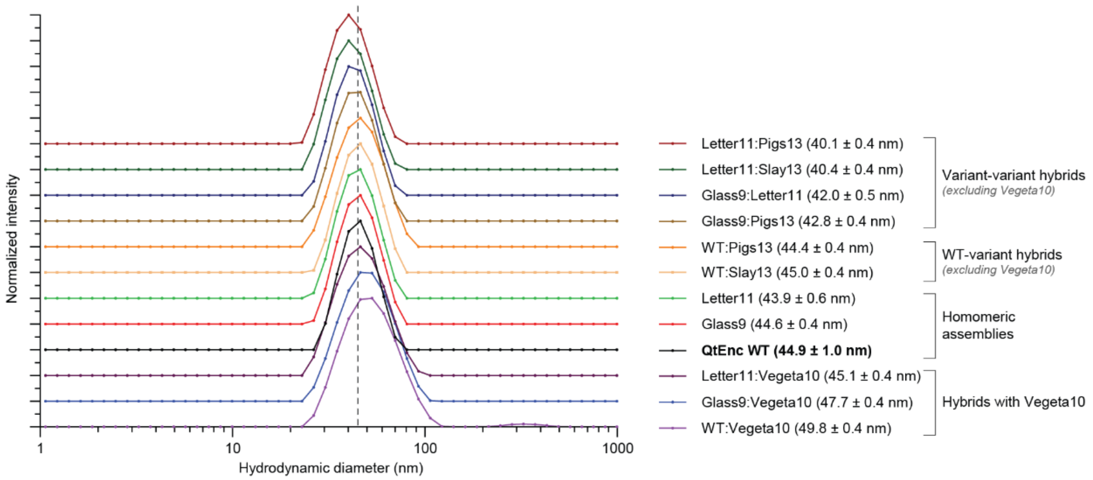


**Fig. S11. Characterization of hybrid assemblies by dynamic light scattering.** All variants and 1:1 hybrids have similar hydrodynamic diameters to QtEnc WT (vertical dashed line indicating WT diameter). Variant-variant hybrid assemblies have slightly smaller hydrodynamic diameters than homomeric assemblies and WT-variant hybrids, with the exception of hybrid assemblies containing Vegeta10 which have a slightly larger average diameter. The error refers to the standard deviation for the cumulant fit over 10 acquisitions. Each data series is displayed with a vertical offset to aid visual clarity.

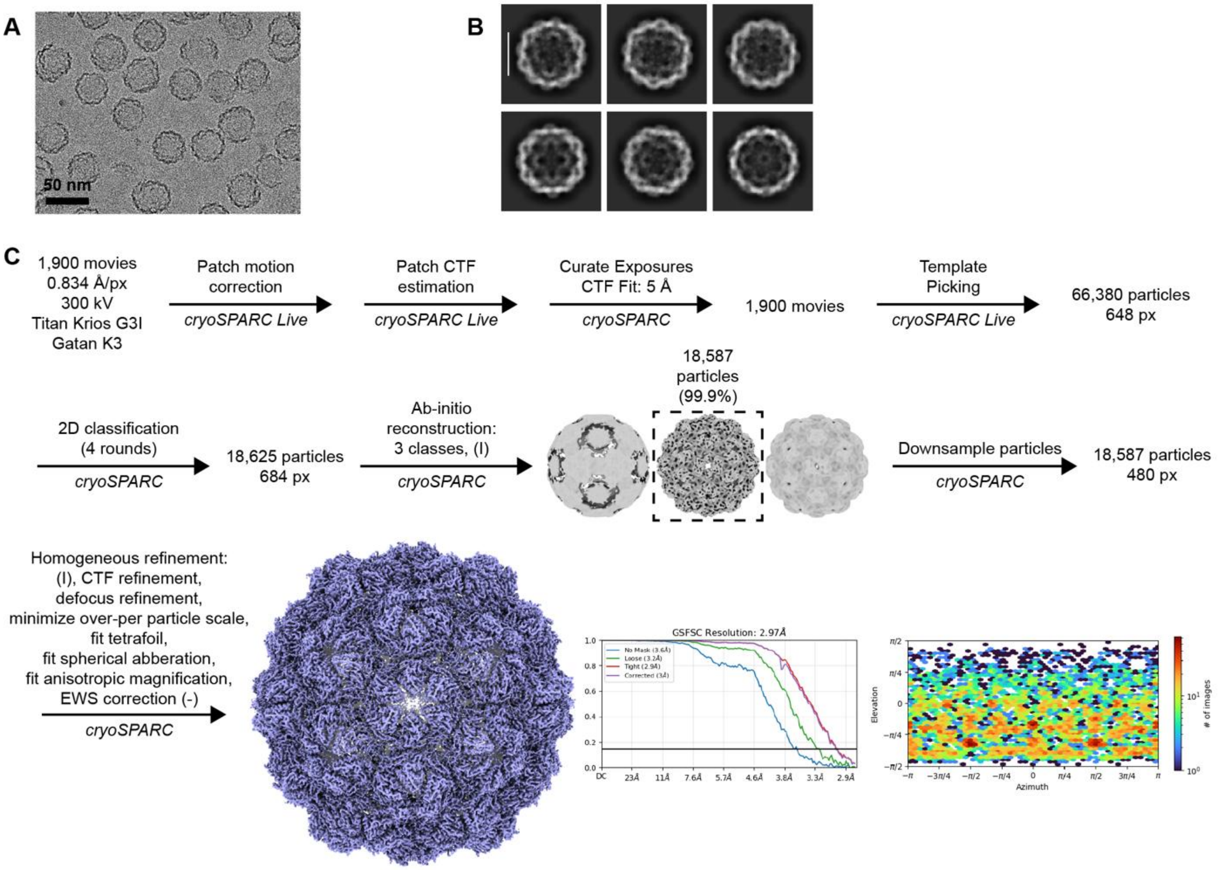


**Fig. S12. Cryo-EM data and processing workflow for the WT:Vegeta10 (1:1) hybrid.** (**A**) Representative raw micrograph. (**B**) Representative 2D class averages (scale bar = 24 nm). (**C**) Cryo-EM data processing workflow.

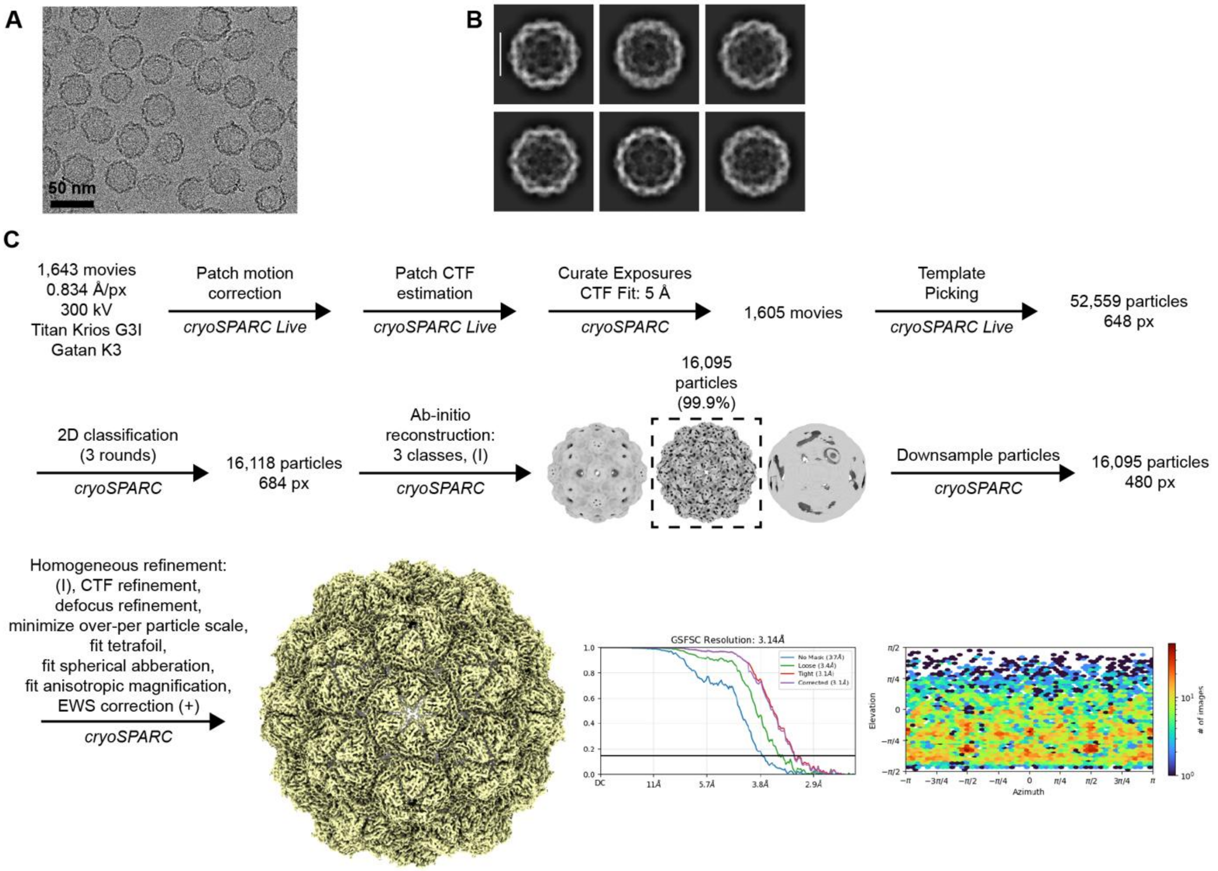


**Fig. S13. Cryo-EM data and processing workflow for the WT:Pigs13 (1:1) hybrid.** (**A**) Representative raw micrograph. (**B**) Representative 2D class averages (scale bar = 24 nm). (**C**) Cryo-EM data processing workflow.

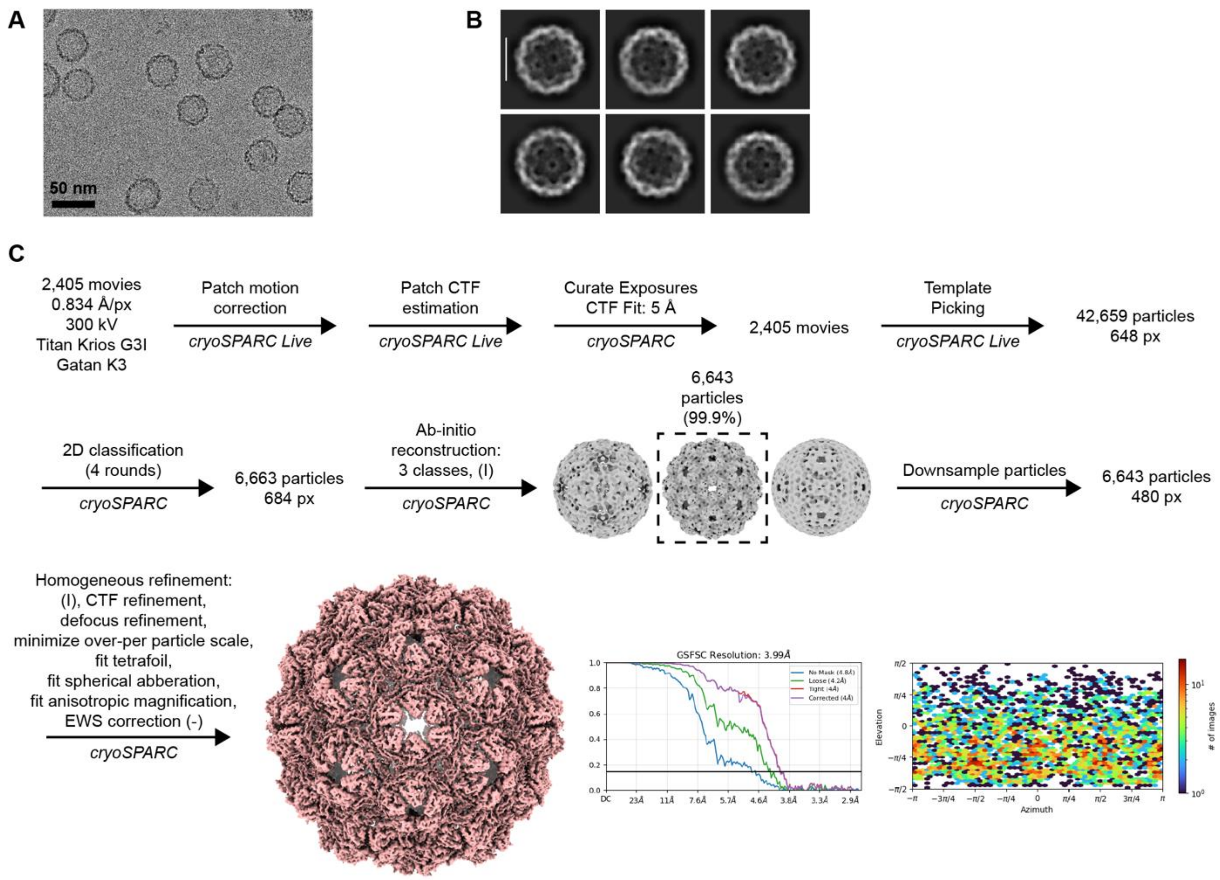


**Fig. S14. Cryo-EM data and processing workflow for the Glass9:Pigs13 (1:1) hybrid.** (**A**) Representative raw micrograph. (**B**) Representative 2D class averages (scale bar = 24 nm). (**C**) Cryo-EM data processing workflow.

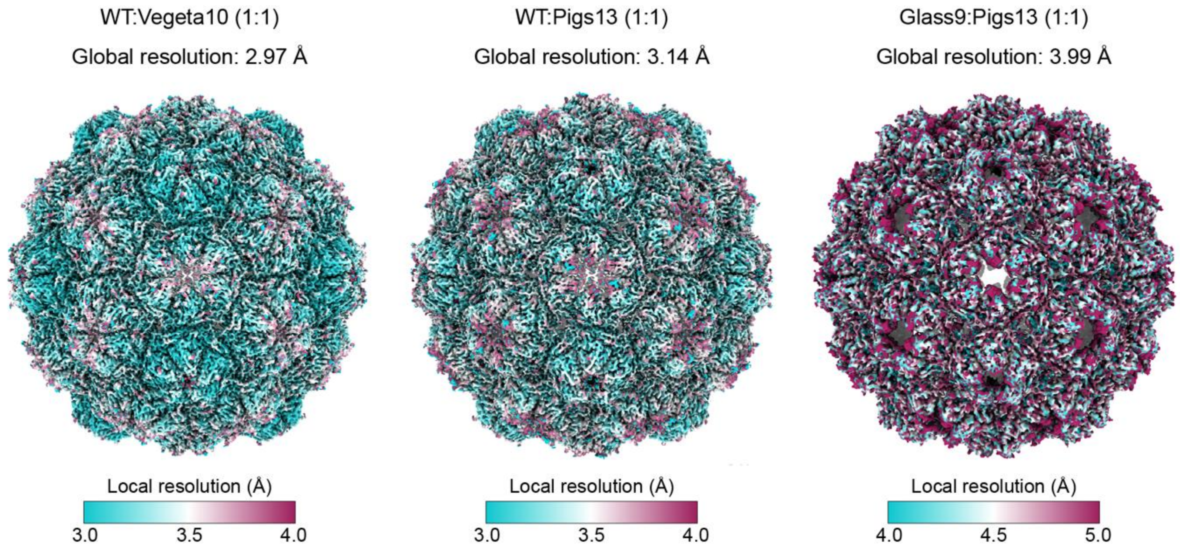


**Fig. S15. Local resolution analysis of the hybrid variants WT:Vegeta10 (1:1), WT:Pigs13 (1:1), and Glass9:Pigs13 (1:1).** Local resolution was calculated in cryoSPARC v4.7.1 at an FSC threshold of 0.5. Global resolution of symmetry-averaged I refinements are shown as well. Maps are colored by local resolution. All hybrids exhibit lower local resolution around the 5-fold (pentameric) and 2-fold (hexameric) pore regions indicating variability. This is consistent with mixed occupancy within stochastically-assembled hybrid shells, as the only differences between protomers are localized at the A-domain pore loops forming the pentameric and hexameric pores. The Glass9:Pigs13 hybrid exhibits large open pores, even at low thresholds, consistent with the fact that both variant protomers that comprise the assembly have a large deletion in the A-domain loop, rather than the loop being disordered and/or missing in the maps.

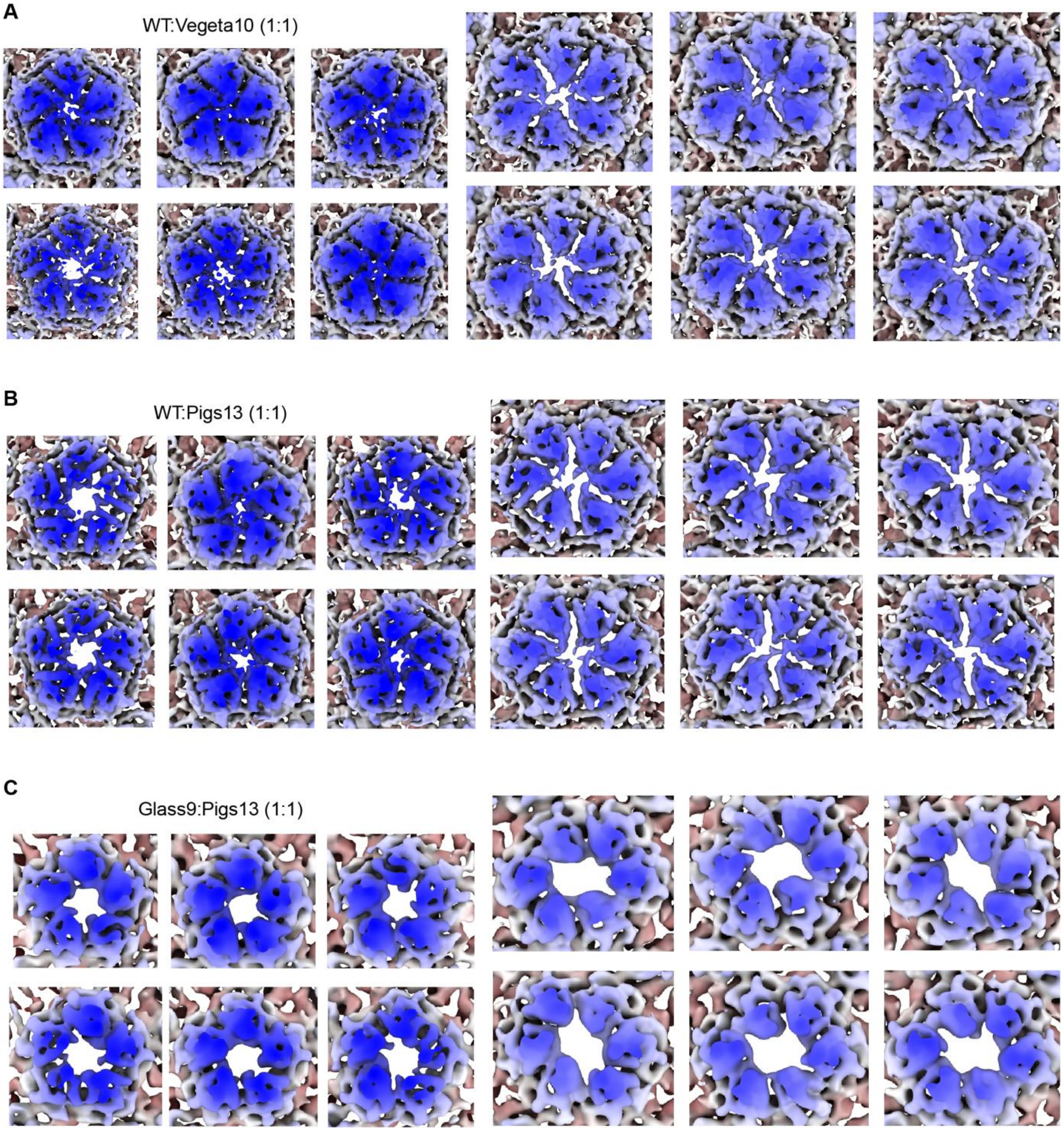


**Fig. S16. Pore analysis of asymmetric C1 cryo-EM reconstructions of the hybrid variants WT:Vegeta10 (1:1), WT:Pigs13 (1:1), and Glass9:Pigs13 (1:1).** Both representative pentameric and hexameric pore regions are shown for WT:Vegeta10 (1:1) (**A**), WT:Pigs13 (1:1) (**B**), and Glass9:Pigs13 (1:1) (**C**). For a given hybrid variant, all pores are shown at the same threshold level. All pore regions show substantial variability consistent with mixed occupancy.

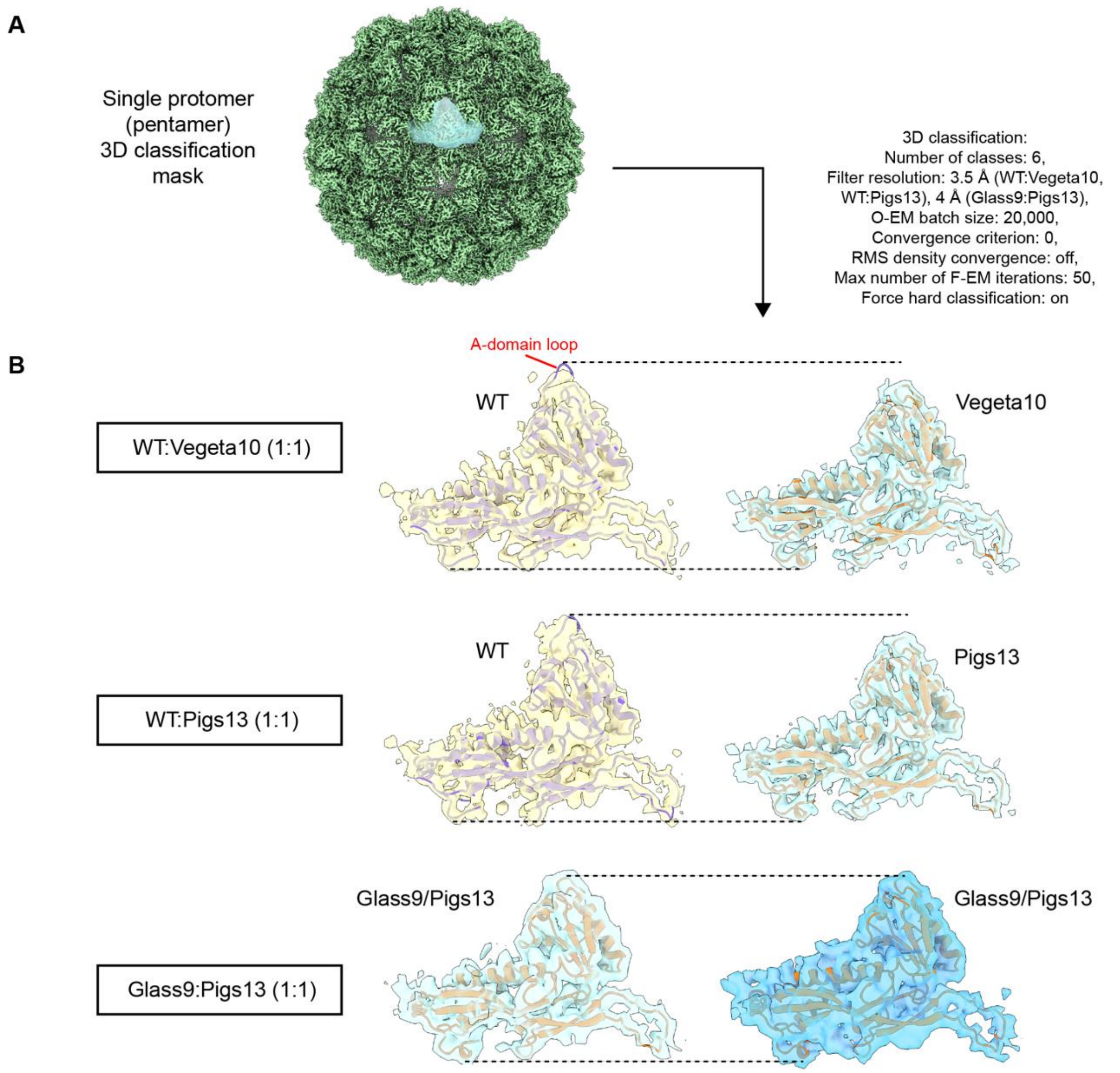


**Fig. S17. 3D classification analysis of the hybrid variants WT:Vegeta10 (1:1), WT:Pigs13 (1:1), and Glass9:Pigs13 (1:1).** (**A**) For 3D classification, a soft mask encompassing a single protomer located at the 5-fold pore was used. 3D classification runs were carried out in cryoSPARC v4.7.1 using the indicated non-standard parameters. (**B**) Representative 3D classes are shown. For WT-containing hybrids, a class resembling WT density (yellow) and a class resembling variant density (blue) are highlighted. The primary difference between WT and variants is the presence of an extended A-domain loop. For the Glass9:Pigs13 hybrid, no classes resembling WT density, containing an extended ordered A-domain pore loop could be identified; therefore, two classes resembling non-WT variants are shown for Glass9:Pigs13. To emphasize density differences/similarities at the A-domain pore loop, all densities are aligned (dotted lines) highlighting that all variants lack A-domain loop density. Within each hybrid, class densities are shown at the same threshold level. WT models (PDB 6NJ8) are fitted into WT-like densities, whereas AlphaFold 3 models are fitted into variant-like densities.

**Table S1-S6 (separate file)**

**Table S1. List of items and reagents**
**Table S2. Oligonucleotide sequences**
**Table S3. Plasmids, descriptions and sequences**
**Table S4. Protein sequences in this study**
**Table S5. Cryo-EM data collection, refinement, and validation statistics**
**Table S6. Uncropped gels and plates**
